# CIRCUST: a novel methodology for temporal order reconstruction of molecular rhythms; validation and application towards a daily rhythm gene expression atlas in humans

**DOI:** 10.1101/2022.12.21.519625

**Authors:** Yolanda Larriba, Ivy C. Mason, Richa Saxena, Frank A.J.L. Scheer, Cristina Rueda

## Abstract

The circadian system drives near-24-h oscillations in behaviors and biological processes. The underlying core molecular clock regulates the expression of other genes, and it has been shown that the expression of more than 50 percent of genes in mammals displays 24-h rhythmic patterns, with the specific genes that cycle varying from one tissue to another. Determining rhythmic gene expression patterns in human tissues sampled as single timepoints has several challenges, including the reconstruction of temporal order of highly noisy data. Previous methodologies have attempted to address these challenges in one or a small number of tissues for which clock gene evolutionary conservation is assumed to be preserved. Here we introduce CIRCUST, a novel CIRCular-robUST methodology for analyzing molecular rhythms, that relies on circular statistics, is robust against noise, and requires fewer assumptions than existing methodologies. Next, we validated the method against two controlled experiments in which sampling times were known, and finally, CIRCUST was applied to 34 tissues from the Genotype-Tissue Expression (GTEx) dataset with the aim towards building a comprehensive daily rhythm gene expression atlas in humans. The validation and application shown here indicate that CIRCUST provides a flexible framework to formulate and solve the issues related to the analysis of molecular rhythms in human tissues. CIRCUST methodology is publicly available at https://github.com/yolandalago/CIRCUST/.

## 1 Introduction

Circadian clocks orchestrate metabolic, endocrine, and behavioral functions. The molecular clock drives tissue-specific rhythms in gene expression [1]. More than ∼ 50% of mammalian genes exhibit daily rhythmic expression patterns, although the specific genes that are rhythmic in one tissue may be non-rhythmic in another, and *vice versa*. Based on these fundamental insights, the importance of biological timing has become increasingly recognized in basic research and medicine, with potential implications for the effectiveness of cancer treatments, heart surgery, and pharmacodynamics [2, 3, 4]. A comprehensive human temporal atlas of 24-h rhythms in gene expression across tissues is therefore of great potential value. Due to the invasive nature, repeat human biopsies are limited to very few tissues, and human gene expression rhythms across tissues rely critically on human postmortem tissue banks [5, 6]. Indeed, human postmortem gene studies are very valuable in circadian biology [7, 8]. However, there are a number of challenges when trying to reconstruct 24-h molecular rhythmicity from postmortem datasets, where each donor only provides one timepoint, including among others, possible uncertainty regarding the actual time of death, postmortem delay and its effect on RNA degradation [9], or inter-individual differences in the alignment of tissue rhythms relative to local clock-time.

The goal of this paper is to describe, validate and apply a method for the estimation of rhythmicity of gene expression given noisy data in order to build a daily rhythm gene expression atlas in humans from postmortem samples. In particular, our interest is focused on the identification and analysis of tissue-specific molecular rhythms and clock genes phase relationships in the human body. Because of imprecisions in estimations of time of death and/or unknown underlying biological times, the first challenge is to estimate temporal order among the samples. This problem is known as the temporal order estimation problem and addressing this problem was the first step in our analysis.

The temporal order estimation problem can be mathematically formulated as that of looking for an *m*-dimensional vector that provides what is known as a *circular order* ***o*** = (*o*_1_, …, *o*_*m*_)^*′*^, where *m* denotes the number of sample collection times to be ordered, see Figures 1 and 2 as illustration. In practice, a circular ordering represents up to 2*m* distinct sample collection time configurations along the 24-h day, depending on the choice of the starting point and the orientation (clockwise or counter-clockwise). The choice of directionality is not trivial and plays a key role in correctly identifying the timing of biological processes across the day, see the Methods section for details.

**Figure 1:**
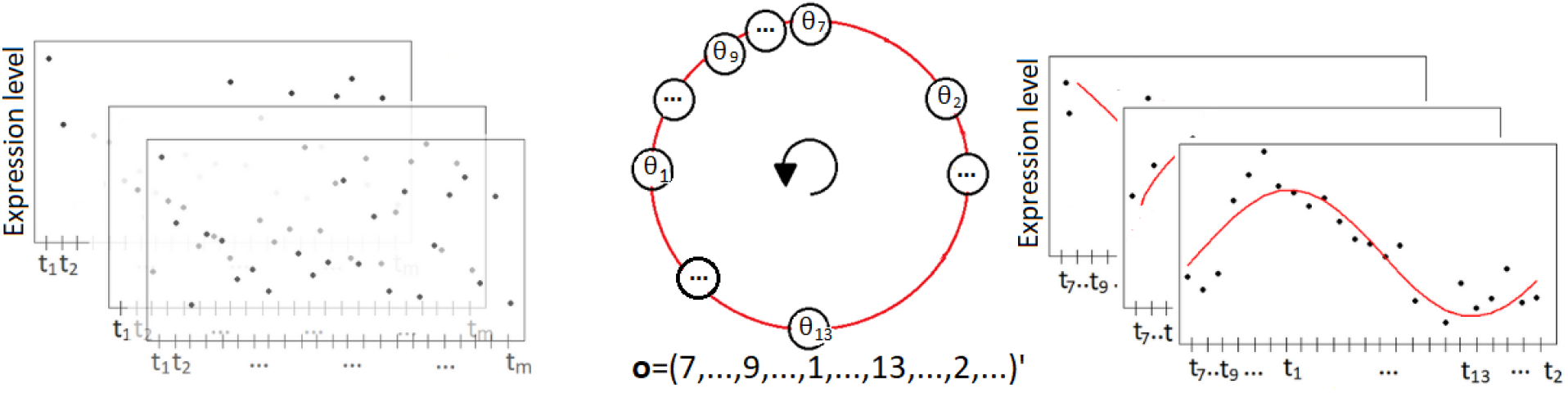
Illustrative outline of CIRCUST solution to temporal order estimation conducted at each tissue. Left: Unordered gene expression data across *m* samples registered at arbitrary clock times *t*_1_, …, *t*_*m*_ along the 24-h day. Superimposed rectangles are different genes of the tissue. Dots are the gene expression data. Middle: Circular order ***o*** obtained from ***θ***, where 0 ⪯ *θ*_7_ ⪯ … ⪯ *θ*_9_ ⪯ … ⪯ *θ*_1_ ⪯ … ⪯ *θ*_13_ ⪯ … ⪯ *θ*_2_ ⪯ 2*π*. Starting point and direction are fixed so the assumptions considered are fulfilled. Right: Ordered gene expression data, as a function of CIRCUST estimated times, across *m* samples registered at clock times *t*_1_, …, *t*_*m*_ along the 24-h day, where *o*_*j*_ = *k* ⇔ *t*_(*j*)_ = *t*_*k*_, ∀*j* = 1, …, *m, k* ∈ {1, …, *m*} and *t*_(*j*)_ is the *j*-th element in the vector of ordered timepoints. Superimposed rectangles are the different genes of the tissue. Dots are gene expression data.

**Figure 2:**
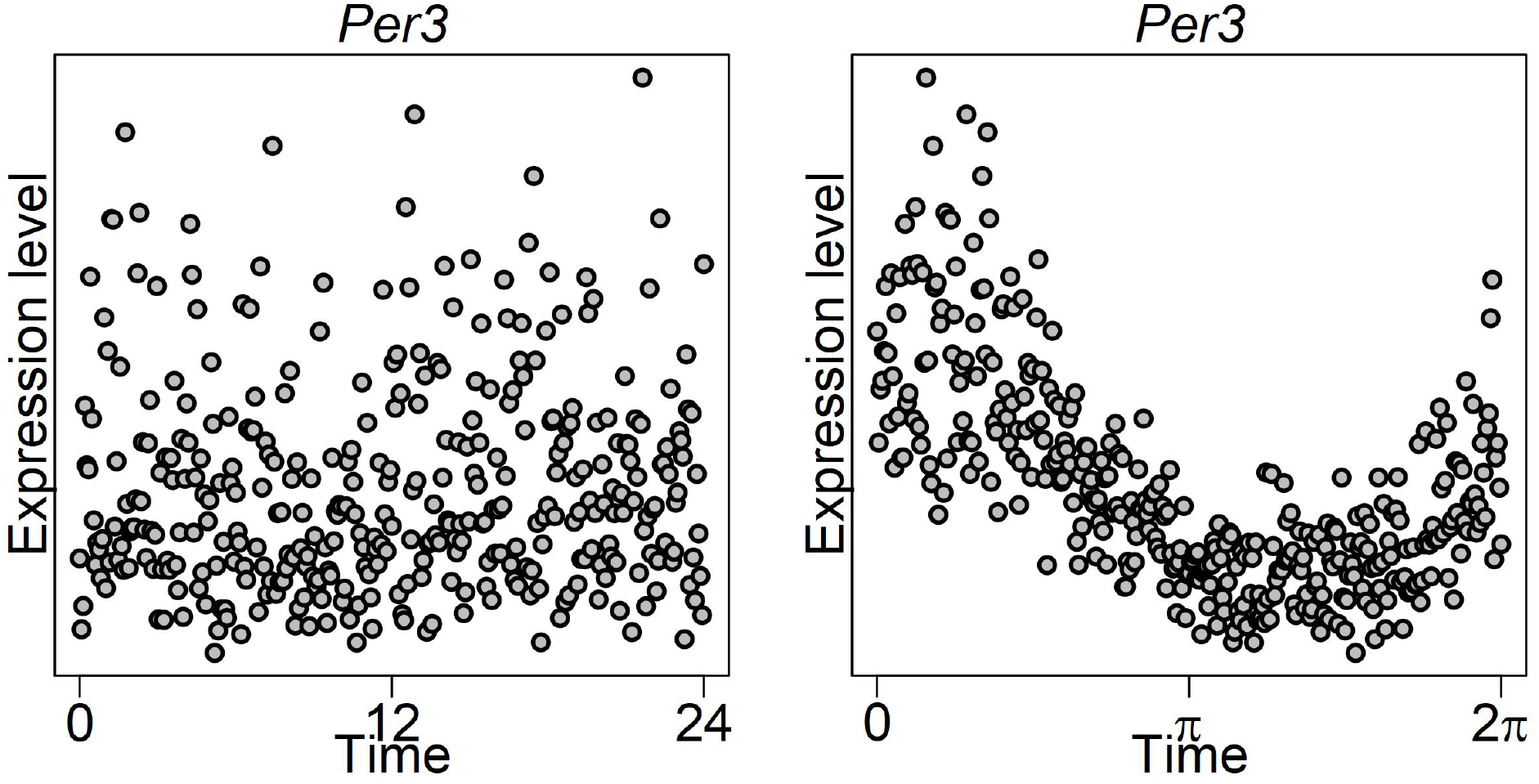
TOD versus CPCA sampling time estimates for gene *Per3* on Skin Not Sun Exposed (Suprapubic) tissue from GTEx. Left: Gene expression as a function of TOD times. Right: Gene expression as a function of CIRCUST estimated times.

This problem has recently garnered a lot of interest within circadian biology, and several methodological approaches have emerged depending on the problem at hand, including Oscope [10], reCAT [11] and CYCLOPS [12] among the most extensively used in practice. Oscope and reCAT were specifically developed to recover cell-cycle dynamics from unsynchronized single-cell transcriptome data, and are highly sensitive to inter-subject variability, as those observed in human gene studies. CYCLOPS, based on a neural network approach overcomes these drawbacks, but it requires evolutionary conservation of the relationship between the rhythmicity of individual clock genes across species which is not always known [13]. Even so, CYCLOPS has been used widely, however only for one or a limited number of tissues [14, 15]. More recently, we have introduced a non-parametric framework to mathematically formulate and efficiently solve the temporal order estimation problem without any additional genomic information, but for the case of equally-spaced timepoints, which is an assumption that is not met in postmortem gene studies [16].

After estimating temporal order, it is needed to identify tissue-specific molecular rhythms, as well as to assess peak phase relationships across tissues. Several models have been proposed in the literature for the analysis of oscillatory rhythms, referred to hereafter as *rhythmicity models*.

Cosinor [17] is the classical rhythmicity model widely utilized in chronobiology [7, 8, 12, 14]. It is a parametric model that consists of three parameters and captures rhythmic patterns using a sinusoid. Yet, Cosinor may be too rigid for the analysis of transcriptome data exhibiting asymmetric patterns (see Figure 3). Cosinor can be extended to a multi-component model by including multiple sinusoidal harmonics to gain flexibility. Even in this case, it may be unsuitable for the analysis of molecular rhythms as was shown in [18]. In addition, the use of a large number of components may result in serious overfitting issues.

**Figure 3:**
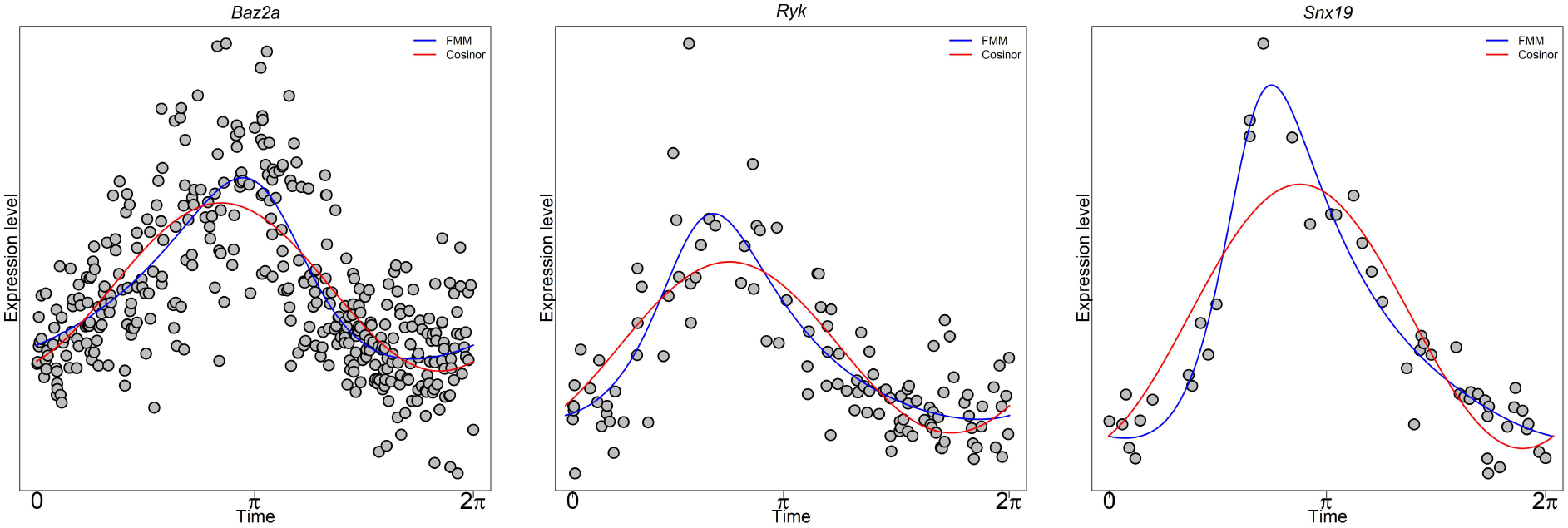
FMM versus Cosinor performance on selected TOP genes from different GTEx tissues. *Baz2a* (left), *Ryk* (middle), and *Snx19* (right) gene expression as a function of CIRCUST estimated times from Lung, Small Intestine, and Kidney, respectively. FMM predictions are shown as blue solid lines. Cosinor predictions are shown as red solid lines.

Alternative rhythmicity models have emerged. In [19], and references therein, models based on ordinary differential equations are proposed to describe circadian clock dynamics. However, the type of equations and model parameters are arbitrary and highly dependent on the process under study. Within a non-parametric perspective and in the context of the Order Restricted Inference, we developed ORI, a computationally efficient and versatile model, that formulates rhythmicity (up-down-up pattern) by using mathematical inequalities covering a wide range of rhythmic patterns [16]. However, comparing rhythmic patterns with this model is not straightforward, as it is for parametric models. To overcome these drawbacks, we presented Frequency Modulated Möbius (FMM) model, a flexible five-parametric model that allows deformations to sinusoidal shape to accommodate commonly seen asymmetries in applications (see Figure 3) [18]. This is because FMM is formulated in terms of the phase, an angular variable that represents the intrinsic rhythmicity of the oscillation that periodically repeats every 24-h. Moreover, FMM model parameters are easy to estimate providing meaningful interpretations. An overview of the FMM model is given in Section 3.1 of the Supplementary Materials.

This work proposes CIRCUST, a general methodology that solves the temporal order estimation problem, as well as identifies and characterizes a wide variety of genes that express 24-h biological rhythmicity, including those with asymmetric expression patterns. The method makes use of the underlying CIRCular structure of the molecular rhythms [16], and the robUSTness of the mathematical procedure to cope with the high noise levels and inter-individual variability that characterize human postmortem gene studies. Specifically, the temporal order reconstruction problem is addressed by a circular dimensionality reduction approach called Circular Principal Component Analysis (CPCA), see Methods section, while the use of the FMM rhythmicity model provides precise estimates of the rhythmicity parameters such as phase.

There is no gold-standard dataset with repeated sampling across multiple human tissues and most human studies have been limited to blood (e.g., [20]) or to another tissue with low sampling frequency [21, 22]. Additionally, inter-individual variability increases uncertainty of estimation of biological timing [23]. This paper shows that CIRCUST is a sound framework based on the analysis of molecular rhythms from two controlled experiments. The first validation dataset consists of human epidermis, a tissue with robust circadian oscillations, repeatedly collected at known and unknown timepoints across a 24-h timeframe from healthy adults [15]. The second validation set consists of a large set of different tissues collected at known timepoints across a 24-h timeframe from baboons, a primate closely related to humans [24]. Next, the Genotype-Tissue Expression (GTEx) dataset, a postmortem gene expression dataset across the largest number of human tissues was considered [25]. GTEx provides annotated times of death (TODs) estimates. However, such TODs may give inaccurate information, see Figures 2 and S1, because of the large inter-individual differences in the timing of the central circadian pacemaker, even in healthy patients [26, 27, 28]. CIRCUST was conducted on GTEx towards developing an atlas of human 24-h expression rhythms across a wide range of tissues that may provide novel insights into the molecular clock networks.

## 2 Methods

The CIRCUST methodology includes reconstruction of temporal order followed by estimation of rhythmic parameters. The details are described below.

### 2.1 CIRCUST solution to temporal order estimation

For each tissue, CIRCUST addresses temporal order reconstruction based on Circular Principal Component Analysis (CPCA), a simple and efficient approach to the sampling time estimation problem. CPCA is a nonlinear dimensionality reduction method that describes the potential circular structure of the molecular rhythms by its projection onto the unit circle [29, 30]. CPCA is often computed from a sub-matrix of a reduced number of tissue-specific rhythmic genes, instead of considering the raw gene expression matrix. Two different sets of rhythmic genes are considered in this paper: a set of 12 well-established core clock genes for an early stage; and subsets of tissue-specific markedly rhythmic genes, called TOP rhythmic genes, at later stages, see below for details.

The CPCA-based solution starts with the computation of the two first *eigengenes* from a sub-matrix of rhythmic gene expressions. Eigengenes are gene-like expression patterns across samples obtained as a linear combination of the expressions in the matrix [31]. Despite the initial unordered expression patterns of these two eigengenes, its mapping reveals the underlying circular structure over the samples, as illustrated in Section 3.2 of the Supplementary Materials. Next, eigengenes are projected onto the unit circle [0, 2*π*), computing the arctan of these projections which allows defining the angular vector ***θ*** = (*θ*_1_, …, *θ*_*m*_)^*′*^ that represents the temporal position of the *m* samples in the raw gene expression sub-matrix onto the unit circle. The increased order of these angles sets the circular order ***o*** = (*o*_1_, …, *o*_*m*_)^*′*^ which provides a circular arrangement of the timepoints. Finally, for the given order, there exist 2*m* sample time configurations according to the starting point and the (clockwise/counterclockwise) direction selection. In general, this choice is made so that two standard assumptions concerning the core clock genes’ peak phase relations in mammals are fulfilled see Section 3.2 in the Supplementary Materials for details). These assumptions can be user-refined, in terms of peak phases’ order restrictions, in case the molecular clock network of the species is (partially) known *prior*, yielding more reliable sampling time estimates. We refer to this particular case as CIRCUST_*prior*_. Full details regarding temporal order estimation are given in Section 3.2 of the Supplementary Material. Figures 1 and 2 illustrate a CPCA solution to approach the temporal order identification problem.

### 2.2 CIRCUST methodology

Let [***R***] denote the matrix of raw and unordered expressions data that serves as input. For each tissue, CIRCUST is sequenced as follows. Figure S2 shows an outline of the methodology.

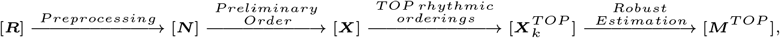

where [***N***] is the matrix of preprocessed, normalized (and unordered) expression data. [***X***] is a preliminary ordered gene expression matrix, and 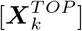 is the *k* − th expression matrix with the ordered expression data of the tissue-specific *TOP* genes, i.e. the highly rhythmical genes of each tissue, *k* = 1, …, *K* with *K* a prefixed integer value (see below). To define these two latter (ordered) matrices the temporal order problem must be addressed. The output of CIRCUST is [***M*** ^*T OP*^], a matrix that contains robust (Median) of the main FMM parameter estimates computed for the *TOP* genes in 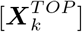, *k* = 1, …, *K*. FMM parameters are meaningfully interpretable and characterize rhythmicity, see Section 3.1 in the Supplementary Materials. CIRCUST steps are described below.

1. ***Preprocessing*** Genes with zero read counts in more than 30% of samples are discarded. Gene expressions are one by one normalized into [-1,1] by using a min-max normalization [16]. The preprocessed expression matrix is denoted by [***N***].
2. ***Preliminary order*** A core information set consisting of the 12 clock genes: *Per1, Per2, Per3, Cry1, Cry2, Arntl, Clock, Nr1d1, Rora, Dbp, Tef* and *Stat3* is considered. There is no a gold-standard for core clock genes selection, though gene expression patterns of this choice, generally display marked circadian signals in most of the human tissues [1] and were also considered as circadian benchmarks in previous works [12, 24, 14, 15]. The role of CPCA at this point is twofold. CPCA is computed on the sub-matrix of the 12 core clock genes from [***N***]. First, CPCA allows detecting outlier samples, see Section 3.3 in the Supplementary Materials for details. Outliers are deleted from [***N***], and the expression data are normalized again. Second, CPCA provides a solution for the temporal order identification problem (setting starting point and direction), from the sub-matrix of the 12 core genes from [***N***], as was detailed above. Then, [***N***] is ordered with regard to the circular order obtained as the solution of CPCA. We refer to this matrix by [***X***].
3. ***TOP rhythmic orderings***. Rhythmicity models are used at this stage to predict gene expression patterns. First, the ORI model’s [16] computational efficiency allows discarding potentially non-rhythmic genes, with 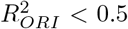, in [***X***]. *R*^2^ is a rhythmicity model’s goodness of fit measure taking values from 0 to 1; the closer to 1, the higher the rhythmicity, see Section 3.4 of the Supplementary Material. Then, the tissue-specific *TOP rhythmic genes* are defined, based on the FMM model predictions, as those which are: i) non-spiked 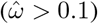; ii) with the highest rhythmicity 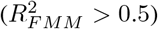; and iii) whose peak phases 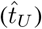cover all the quarters of the unit circle ([0, 2*π*)). This definition results from the meaningful interpretation of the FMM parameters: *ω, t*_*U*_, see Section 3.1 in the Supplementary Materials and [18] for details. The 12 core clock genes are usually among the TOP genes, if not, they are forced to be included. [***X***^*T OP*^] denotes the sub-matrix of TOP genes once they are filtered from [***X***]. Next, random selections of size 2*/*3 of the genes in the TOP are considered. CPCA-based solution for temporal order estimates is recomputed for each of these sub-matrices resulting from filtering the selected genes of [***X***^*T OP*^]. The process is repeated until obtaining a prefixed number of *K* random gene collections verifying that: (a) angular values in ***θ*** are distributed along with more than half of the unit circle; (b) and the maximum distance between two consecutive angular values in ***θ***, does not exceed the observed distances for any pair of consecutive angular values with regard to the preliminary order given by the vector ***θ*** considered in step 2. Hence, ***o***_*k*_, *k* = 1, …, *K* circular orders are defined. For each of them, [***X***^*T OP*^] is reordered, obtaining 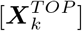, that denotes the *k*-th matrix of TOP genes ordered by ***o***_*k*_, *k* = 1, …, *K*.
4. ***Robust Estimation***.

FMM predictions for the TOP genes in 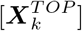, *k* = 1, …, *K*, are computed. For each gene at the TOP, there are *K* FMM parameter estimates, and *K* rhythmicity measures 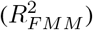. Robust FMM parameter estimates, in terms of the medians, are computed. [***M*** ^*T OP*^] is the matrix that contains for the genes in the TOP the median of the FMM features: *R*^2^, *t*_*U*_ and *ω* which are key to assess and compare rhythmicity across tissues.

## 3 Results

In this section, CIRCUST is validated using ordered data from humans and baboons. We also illustrate the application of CIRCUST to GTEx towards developing a daily rhythm gene expression atlas in humans.

### 3.1 CIRCUST validation on human epidermis

This validation relies on the hybrid human gene expression dataset from epidermis tissue (GEO accession number GSE139301) [32]. On the one hand, this dataset contains gene expressions for a set of 19 participants for which biopsies were collected at 6 am, 12 pm, 6 pm, and 12 am. On the other hand, it includes the gene expression for 533 epidermis samples for which sample collection times were unrecorded.

We apply CIRCUST on the set of 533 unordered samples in order to compare the results with those obtained for the 19 participants where clock times were known. This latter mimics what was done in [32] to validate CYCLOPS. For comparison purposes, the analyses refer to the set of clock-associated genes in [32] which are among those at the TOP genes of CIRCUST for epidermis tissue see Figure 4. Gene expression data from biopsies at the four timepoints for the 19 participants are displayed in Figure 4 (A), see thin color lines. Due to the low sampling frequency and the noise inherent in the experiment, for each gene, the averaged expression pattern is computed (blue thick line). FMM model predictions for the average expression patterns are computed, assuming that the estimated FMM peaks vas the peak phase values for these genes. Figure 4 (B) compares these peak values (triangles) derived from the 19 participants, where clock times are known, with the estimated peak phases derived from CIRCUST (circles) and CYCLOPS (squares) for the 533 samples, when sampling times are unknown. The circular correlation [33] between participant peak phases and estimated phases from CIRCUST and CYCLOPS for these genes are 0.862 and 0.819, respectively, revealing coherence between the CIRCUST peak phases’ estimates and the peak phases given for known clock times. In particular, the differences between CIRCUST and peak phases from the participants are, in general, less than ∼ 2 hours (∼ 0.52 radians), being especially low for the genes *Per2, Cry1, Cry2* and *Ddp*. Except for *Per3*, such differences tend to be lower for the CIRCUST than for the CYCLOPS. Moreover, the orders among peak phases defined by CIRCUST match with those observed from the human biopsies across the 19 participants: {*Per3, Dbp, Tef, Ciart*} ⪯ {*Cry2, Per2*} ⪯ {*Cry1, Arntl*} ⪯ {*Per3, Dbp, Tef, Ciart*}, ⪯ is read as “before than”. Finally, the rhythmicity measures 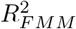 for the eight TOP genes (see Figure 4 (C)) appear to be more consistent with the oscillatory expression patterns observed in Figure 4 (A) than those given by CYCLOPS in [32], with *Ciart, Tef* or *Arntl* being among those that display the strongest rhythmicity.

**Figure 4:**
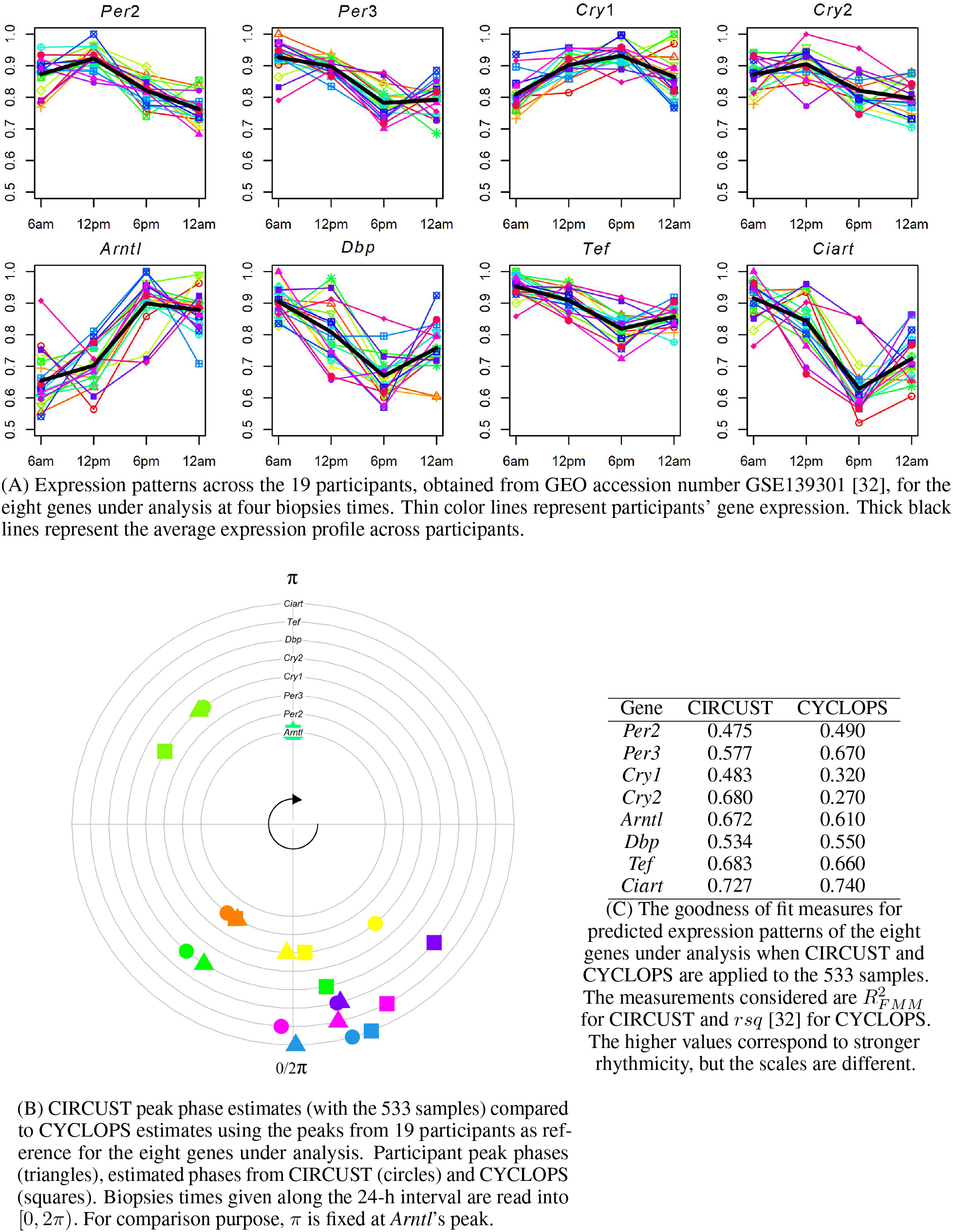
CIRCUST consistency for human epidermis (GSE139301) dataset

### 3.2 CIRCUST validation on multiple baboons tissues

The second validation is driven by the baboon gene expressions dataset (GEO accession number GSE98965). Data were collected, under controlled conditions, every 2 hours (ZT0, ZT2,…, ZT22) over the 24-h day across 64 different tissues, which are aggregated into 13 functional groups [24]. In order to guarantee the consistency of the results, analyses are restricted to the 47 baboons’ tissues for which the rhythmicity measure 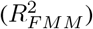 for the 12 core clock genes is, on average, higher than 0.7, see Table S1 for details. Among these tissues, there are representatives of 12 out of 13 of the functional groups, all except for the male genitals. The baboon is a well-studied mammalian species in circadian biology with well-established prior knowledge regarding its molecular clock network. *CIRCUST*_*prior*_ allows incorporating such information into the method in terms of order peak relationships (inequalities) improving its performance (see Methods section). Specifically, [1, 12, 34, 35] reported that baboons’ peak phases usually fulfil: {*Dbp*} ⪯ {*Cry1, Cry2*} ⪯ {*Arntl*} or {*Nrd1*} ⪯ {*Per1, Per2, Per3*} ⪯ {*Arntl*}. In case one of the relationships above increases the number of core clock genes with their peaks within the active period ([0, *π*)) with regard to the standard order peak time assumption (2) (see Subsection 2.2 in the Supplementary Materials), it will be replaced by the specific relation given for baboons.

Circular association between CIRCUST estimated times in [0, 2*π*) and the real times (ZT0, ZT2,…, ZT22) along the periodic scale of 24-h, which can be represented as points on a circle, is assessed. Both variables can be considered as angular, then a circular-circular regression problem [36], similar to the linear regression when both variables are euclidean, is solved. For each tissue, the goodness of fit measure *ρ*, defined as an analog of residual sums of squares in a linear regression model, is computed to assess the coherence among both orders [37, 38]. A closer *ρ* to 1 indicates a better correspondence between the orders. *CIRCUST*_*prior*_ performs well in ordering the samples across the 47 tissues, see Figure S3. The interquartile boundary (*P*_25_, *P*_75_) for the values of *ρ* across the 47 baboons’ tissues is: (*P*_25_, *P*_75_) = (0.729, 0.895), see Table S1 for details. The estimated order is very close to the real temporal order for highly rhythmic organs such as White Adipose (*ρ* = 0.964); Pancreas (*ρ* = 0.960); Colon (*ρ* = 0.959); or Skin (*ρ* = 0.953), see Figure 5 (A). In addition, Figures 5 (B), S4, S5, and S6 reveal, that CIRCUST conserves rhythmicity across selected clock genes for the four tissues mentioned above. From mere visual inspection, the gene expression patterns in the baboons at times ZT0, ZT2,…ZT22, (top panels of Figure 5 (B)) are closely tracked by expressions obtained as a function of CIRCUST estimated times for these same genes (bottom panels of Figure 5 (B)). Moreover, these plots show that the FMM model accommodates a wide variety of rhythmic patterns with high (close to 1) and similar rhythmicity strength values quantified by 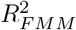, across the selected clock genes, even in those with non-sinusoidal gene pattern, e.g. *Npas2* in Figure 5 (B) and more in Figures S4, S5, and S6.

**Figure 5:**
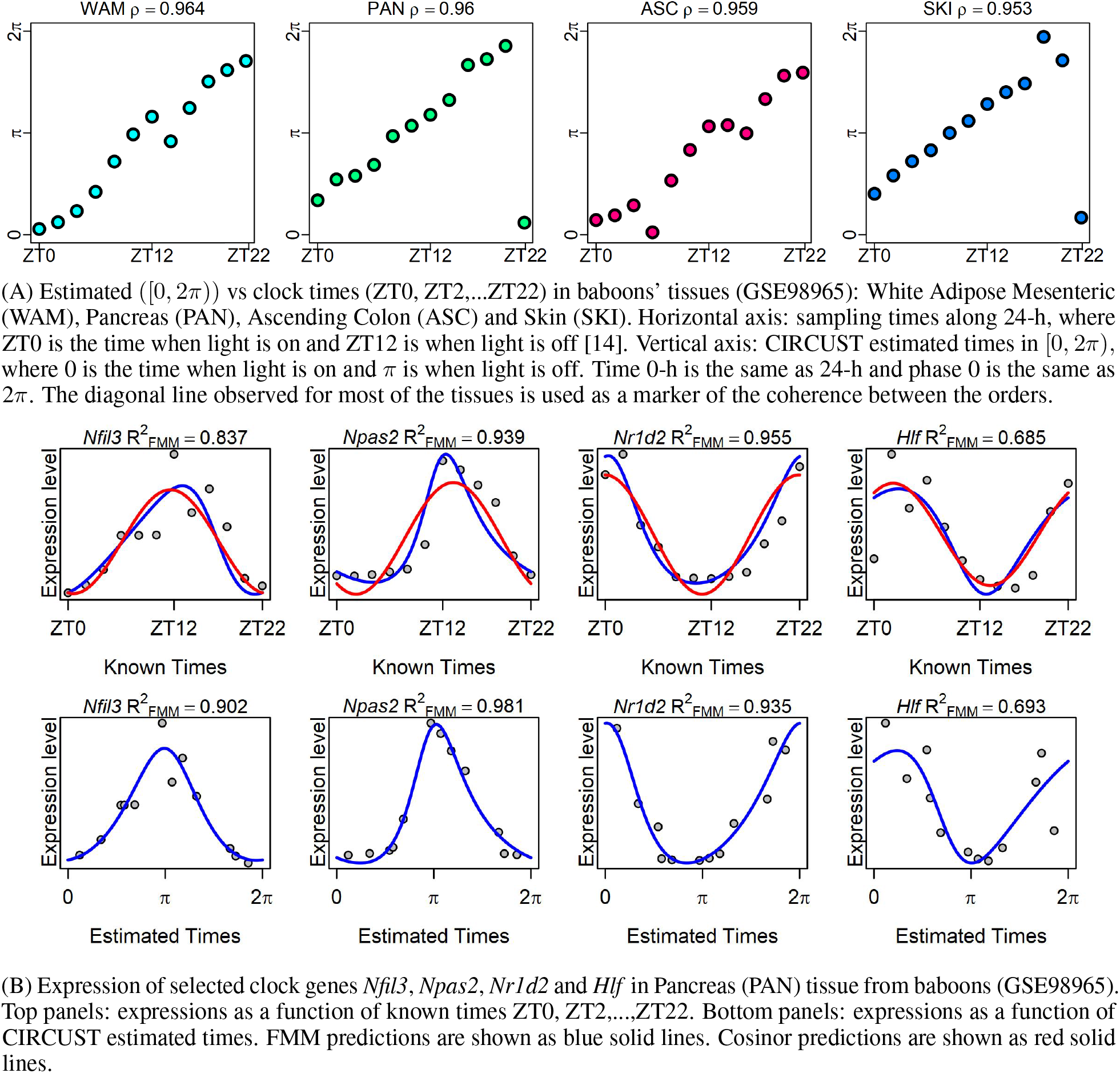
CIRCUST validation based on baboon dataset (GSE98965).

### 3.3 CIRCUST application to GTEx

This section reports the analysis of the molecular rhythms and clock network from GTEx (V7) database. Only tissues with more than 40 samples were included in the analysis. In addition, two cell lines and thirteen brain tissues were discarded [25, 39]. Cell lines may not capture the molecular complexity of the tissue [40]; the brain tissues usually evince intra-tissue heterogeneity and they are often considered as independent molecular networks [41, 42]. Hence, the CIRCUST methodology was separately applied to 34 tissues with a fixed number *K* = 5 of random selections given from the genes at the TOP for each of the tissues. With this criteria, our analysis considers 621 donors characterized by a mix of ages, sex, and health status (see Table S2). Specifically, the results below report, for each tissue, the analyses of the medians of the FMM estimated parameters (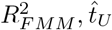, and 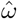) of the TOP genes obtained as outputs (at Step 4) from CIRCUST, see the Methods section for details.

#### 3.3.1 GTEx Molecular rhythm analysis

The molecular rhythms for the TOP genes in each of the 34 tissues from GTEx were analyzed. TOP genes, defined in the Methods section, display non-spike, and heterogeneous rhythmic patterns, as is seen in Figure 3. The number of TOP genes varies among the analyzed tissues (see Figure S7). Muscle-Skeletal, Testis, and Lung are among the tissues with the highest number of TOP genes; while Pancreas or Thyroid are among those with a lower number of them.

Moreover, most of the TOP genes belong to non-intersecting sets (see Figure S8). In particular, for Artery-Tibial and Nerve-Tibial, which are the tissues with the highest number of TOP genes, there are only 5.32% (5 out of 94) shared between both tissues, apart from the 12 core clock genes considered. Moreover, in other rhythmic organs like Testis, 81.61% (71 out of 87) of the genes at the TOP are exclusively rhythmic of this tissue. These latter findings evince tissue-specific rhythmicity in human gene studies.

The heterogeneity observed regarding rhythmicity persists even for core clock genes. Figure 6 illustrates 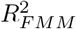 distribution for the genes in the TOP of the 34 organs analyzed. The 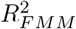 of the 12 core clock genes are shown as different coloured dots. As seen, core clock genes do not always rank among the most highly rhythmic genes of the tissue. Even when analyzing highly rhythmic organs, several scenarios are shown. For example, in Kidney-Cortex, most of the core clock genes are distributed among the TOP genes. On the contrary, the core clock genes in Whole-Blood are not among the TOP genes of this tissue. This latter does not mean that the core clock genes are not rhythmic, but that there are other circadian genes among those in the TOP that, regarding the tissue-specific variability, present a stronger rhythmic signature. In general, for the vast majority of the tissues, at least a quarter of the core clock gene expression oscillations persist across wide inter-individual variability, with clock genes such *Per3* being among those with the highest rhythmicity for more than the 75% of the tissues. This observation supports CIRCUST’s potential to detect novel tissue-specific molecular rhythms in humans, such as *Snx19* in the Kidney, see Figure 3.

**Figure 6:**
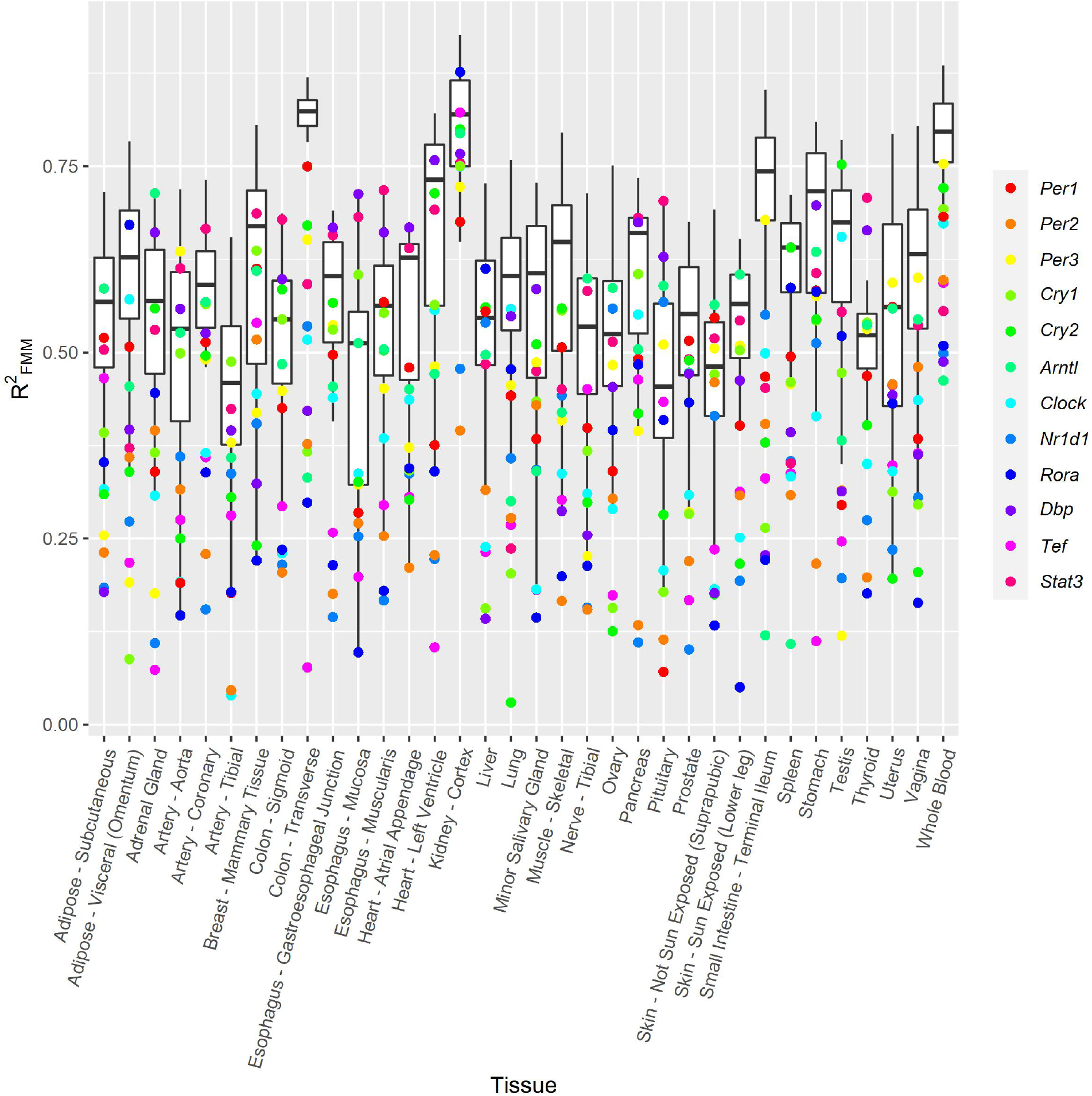
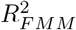 distribution across the 34 organs in GTEx. Dots denote the 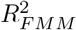 of the 12-core clock. Each core clock gene is represented using a different color. Tissues are alphabetically sorted.

Finally, the atlas of robust human molecular rhythms for the 34 human tissues is provided as a Supplementary data file. For each tissue, the atlas includes the list of TOP genes, ranked from the highest to the lowest rhythmicity, based on the rhythmicity measure 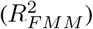, the estimated amplitude (*Â*), peak phase 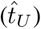, and the timing of the peak phase relative to *Arntl*: corresponding to active/lightened, if 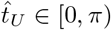, or to inactive/darkness 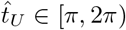. All of these values are derived from Step 4 of CIRCUST methodology. This is the largest rhythmic gene characterization across human tissues to date. This work represents important advances towards a human rhythmic gene expression atlas.

#### 3.3.2 GTEx Molecular clock networks

This section describes and compares the molecular clock networks across the 34 human tissues. The estimated peak phases 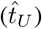 of the TOP rhythmic genes are assessed and compared across the tissues. Here we present data on molecular clock networks simultaneously analyzed across the largest set of human tissues [12, 14, 15].

Figure 7 shows the peak phase distributions of the 12 core clock genes across the 34 tissues. Non-rhythmic core clock genes were discarded from this analysis, see Tables S3 and S4. Distributions varied across organs, but they were not randomly distributed throughout the 24-h day. The peak phase estimates are generally in one or two clusters, with one of them usually preceding the presumed inactive/darkness phase in mammals ([*π*, 2*π*)) [14]. For core clock genes such as *Clock* or *Per1*, peak phase distribution was mainly restricted to a ∼ 6-hour interval. However, for most of the other core clock components, the peaks were distributed along ∼ 12-hour, matching with the presumed active period ([0, *π*)) or light day hours. This reveals human inter-tissue variability and the heterogeneous behavior of the molecular clock networks across the variety of organs analyzed. But, for the particular case of highly rhythmic organs such as Skin (epidermis), molecular clock networks are maintained across species, as shown in Figure 8. There, the estimated core clock peak phases in the skin for humans, from GTEx database, were similar to those estimated for the baboons and both are close to those obtained as a function of the true clock times.

**Figure 7:**
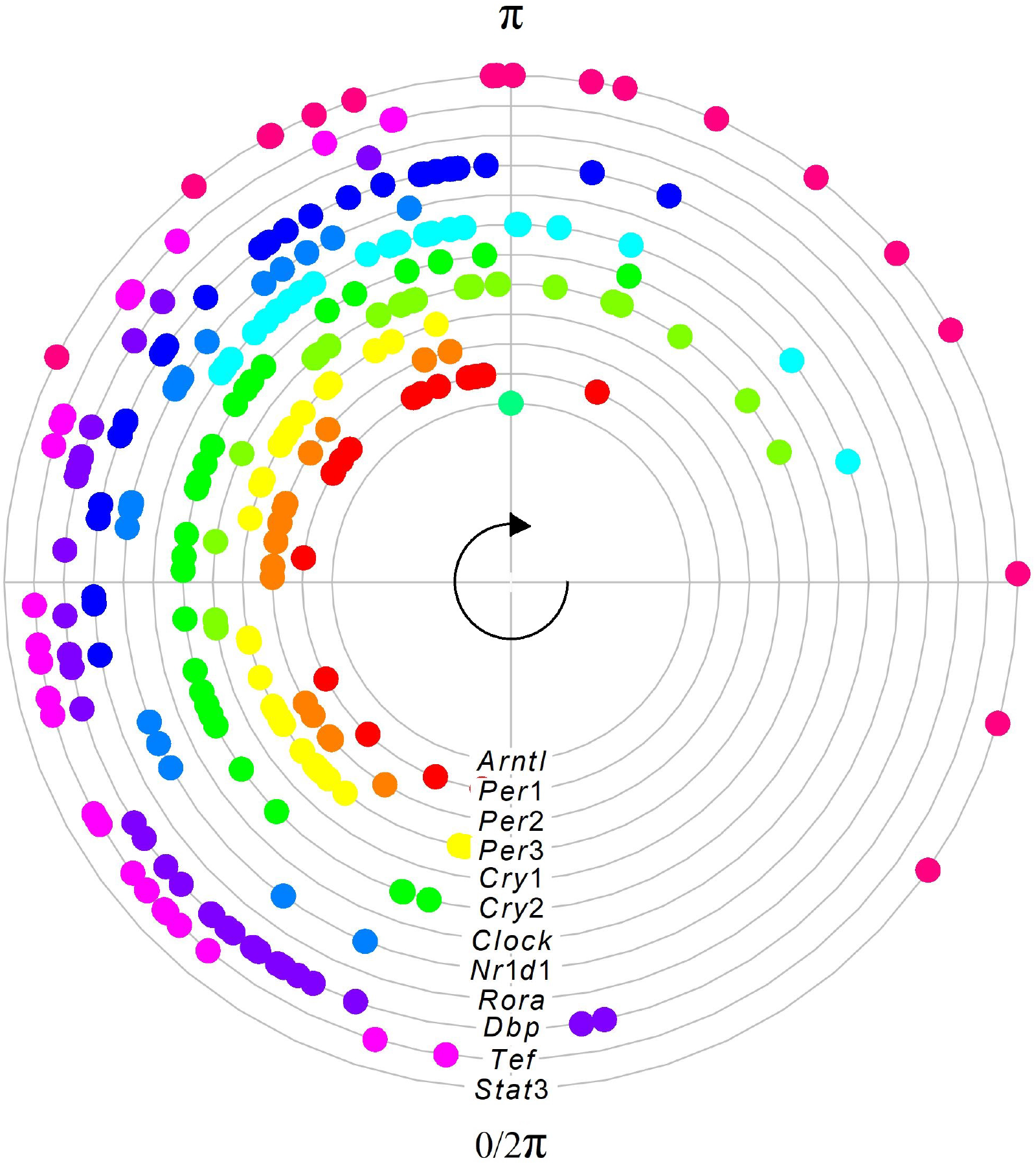
Peak phase estimate distributions of the 12 rhythmic core clock genes across the 34 tissues in GTEx. *Arntl* is known to peak in anticipation of the inactive/darkness [*π*, 2*π*) period in mammals and was set to *π* for comparisons [14]. The *R*^2^ and *t*_*U*_ estimated values are given in Tables S3 and S4, respectively.

**Figure 8:**
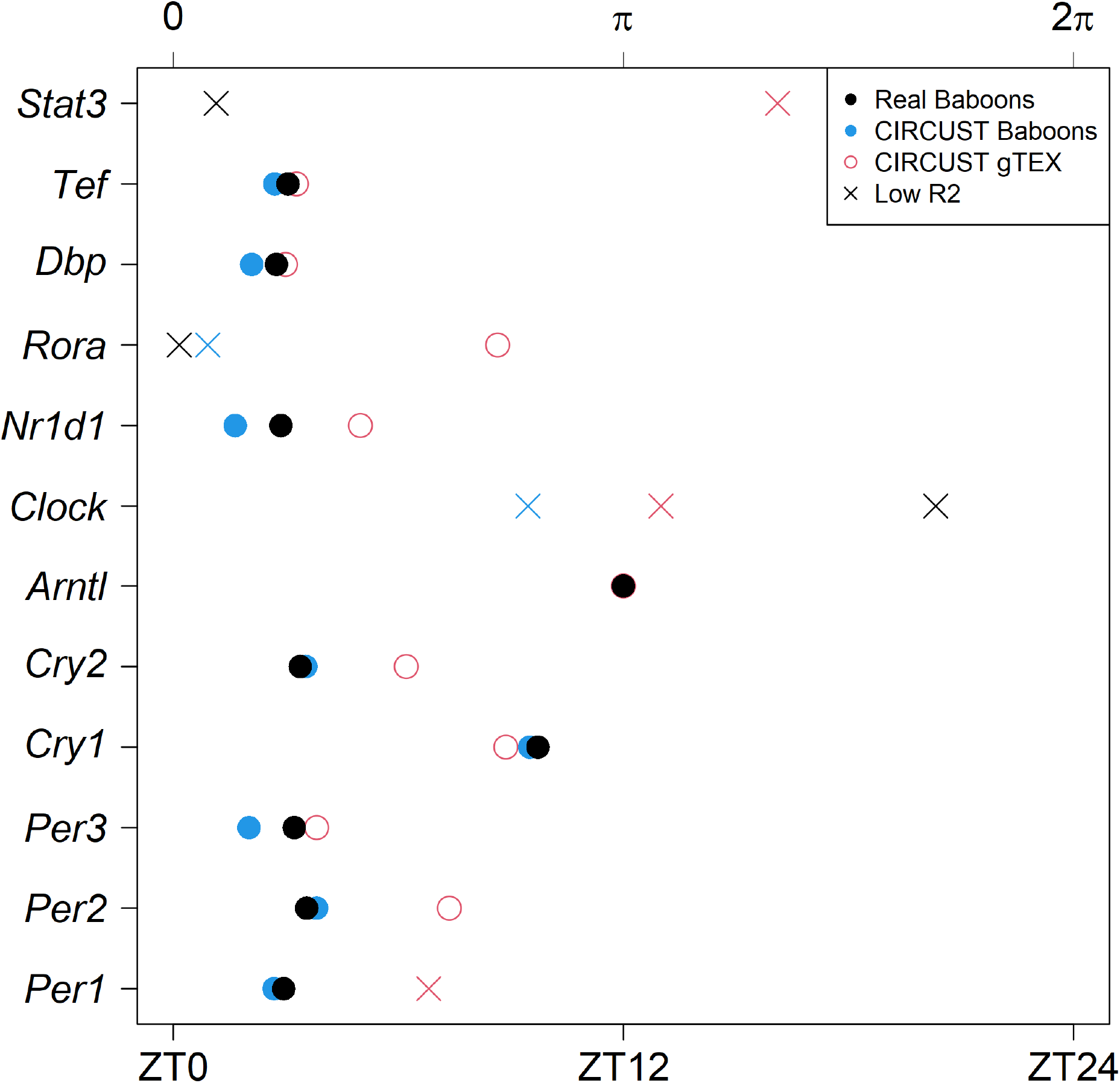
Core clock peaks in the human epidermis (Tissue: Skin - Sun exposed from GTEx) and baboon epidermis (Tissue: SKI from GSE98965) tissue. Estimated phases are derived from the FMM model. Black and blue dots match with the peaks obtained as a function of the true clock times and as a function of the CIRCUST estimated times in baboons, respectively. Red dots match with the peaks obtained as a function of the CIRCUST estimated times in the human epidermis GTEx dataset. Non-rhythmic genes (*R*^2^ *<* 0.5) are marked with a cross.

Finally, the distributions of the peak phases of the TOP rhythmic genes across the human tissues were explored. TOP peaks estimates for nearly all organs display different distributions with one, two, or even three-phase clusters, see Figure 9. In tissues such as Artery-Tibial, Heart tissues, Pancreas, or Stomach, most of the TOP genes peaked within a narrow interval, whereas TOP genes in Colon-Transverse, Spleen, Small Intestine-Terminal Ileum, or Whole-Blood peaked within two distinct time intervals. Three modes are displayed in the Vagina or Testis. Despite human inter-tissue variability, anatomically adjacent tissues showed phased clusters that are temporally close, see for example Esophagus-Gastroesophageal Junction and Colon-Sigmoid, both of which belong to the digestive tract. A compilation of phases across the TOP rhythmic genes revealed that, for the vast majority of tissues, presumed early afternoon major peak anticipating the inactive phase, and a quiescent zone is also observed for many of the tissues which are considered distinctive features of rhythmicity in the diurnal primate [24].

**Figure 9:**
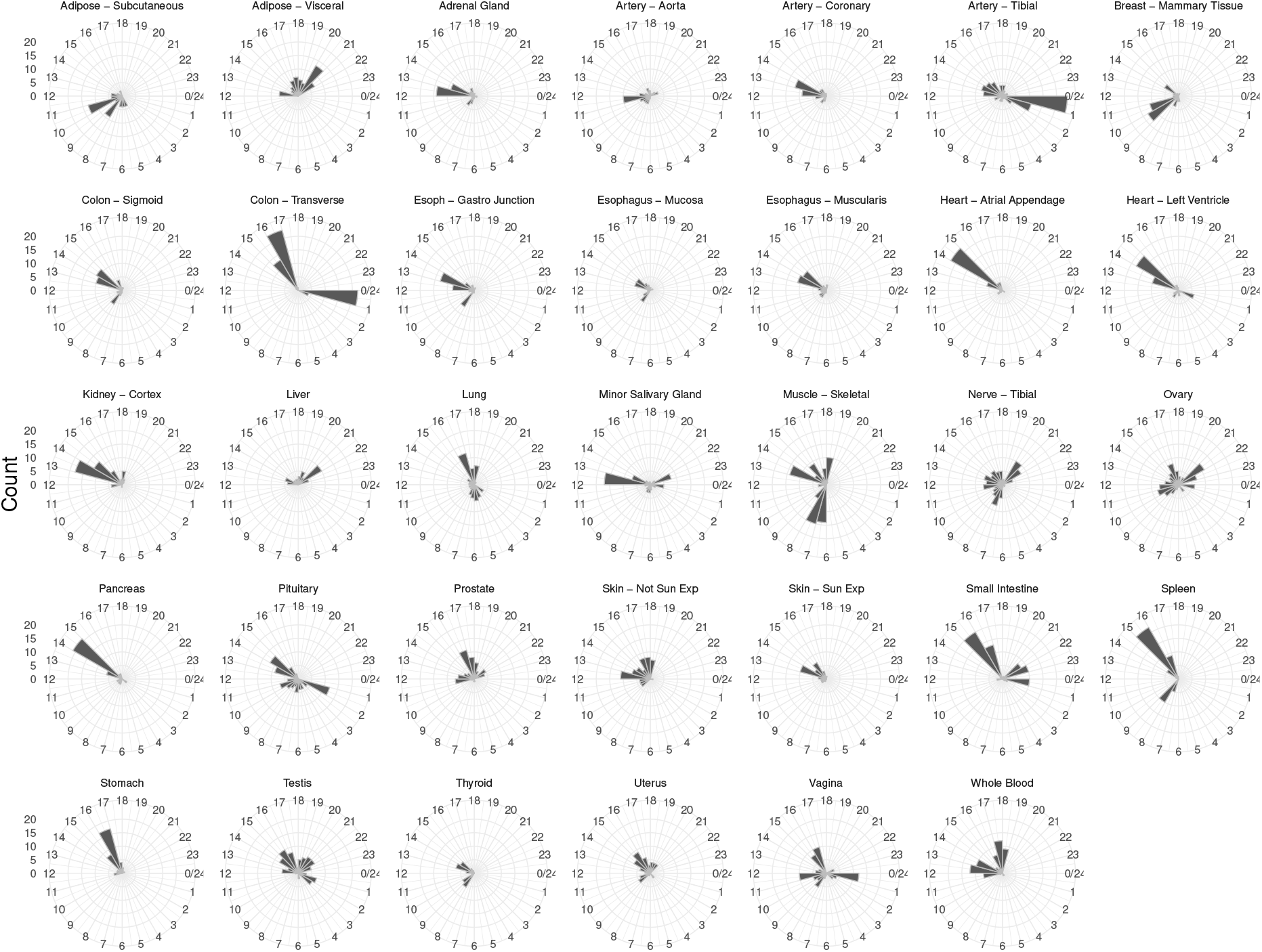
Radial plot of the distribution of the CIRCUST estimated peak phases of the TOP genes along the 34 tissues analyzed from GTEx. Active/light period (0, *π*] is identified with [6am,6pm) and the inactive/dark period (*π*, 2*π*] is done with [6pm,6am) with *π* corresponding with 6pm.

## 4. Discussion

CIRCUST methodology presented in this paper efficiently formulates and solves, based on circular statistics, the temporal order estimation problem arising in gene studies for which the clock time of sample collection is unknown or imprecise. The robustness of CIRCUST against the characteristic noise of postmortem gene studies together with the flexibility of the rhythmicity model FMM expands our knowledge regarding the rhythmicity of genes within and across tissues (see Figure 3). These strengths of methodology give rise to the building of a comprehensive daily rhythm gene expression atlas in humans from the GTEx database that represents rhythmicity analysis across the largest number of human tissues to date.

CIRCUST has a number of advantages compared to existing order reconstruction models [12, 14, 15]. First, CIRCUST does not rely on a black-box model based on machine learning, which allows for a more transparent assessment of the influence of outlier samples. Second, CIRCUST requires only limited assumptions of the temporal order of the core clock or other genes, in that for solving the directionality problem (clockwise or counterclockwise choice), only one comparison between two clock genes is required in the base model (i.e., *Arntl* and *Dbp*). Moreover, not requiring assumptions of the order of rhythmic genes is beneficial because they may not be conserved between model species and humans. This also permits the assessment of a larger number of human tissues as well as application to more diverse species. For example, in [14] only 13 tissues out of the 51 from the GTEx collection were analyzed, in part because in these tissues the phase relationships between the core circadian genes were evolutionarily conserved. This contrasts with the 34 tissues considered in this paper. Beyond the base model, CIRCUST adaptability enables the incorporation of known peak phase relationships between other genes to further enhance the precision of the model.

CIRCUST application can be extended to any tissue of mammal species, regardless of whether or not the molecular clock network is known. Moreover, CIRCUST can be easily adapted to obtain sub-atlases across age, sex, or other variables. To enable wider use, the methodology is open source and publicly available to the scientific community on GitHub https://github.com/yolandalago/CIRCUST/.

It is worth noting that comparison with prior works is challenging as molecular rhythms and clocks network analyses differ across the methods employed, the covariates considered, and the tissues or species studied. Still, we found consistency regarding the tissues with the higher (Artery tibial) and lower (Liver) number of rhythmic genes, as well as the tissues with the higher number of intersecting rhythmic genes (Artery Tibial and Nerve Tibial) with those reported in [14]. Moreover, even though the species considered were different, CIRCUST molecular rhythms analyses in the human GTEx dataset share certain similarities with the results in baboons (a closely related species), where sample collection times were known. In line with that stated in [24], the core clock genes detected by CIRCUST do not systematically rank among the TOP rhythmic genes. This latter finding suggests that core clock gene expression patterns are tissue-specific and their rhythmicity depends on the variability of the tissue and on the rhythmicity strength of the rest of the genes in the TOP. The number of rhythmic TOP genes is relatively lower than in previous works, mainly because of the more stringent requirements of the CIRCUST model. These findings are in line with [42] which reports that the specific genes expressing rhythmicity, including core clock genes, are very tissue-specific. In addition to that, and similar to that described in [24], we discovered that well-established circadian-associated transcripts, such as the recently described *Ciart*, are among the TOP rhythmic genes in more than one-third of the analyzed tissues.

There are substantial differences in the observed daily rhythms in gene expression in the current work compared among species. For example, as observed here, the ranking of highly rhythmic tissues previously documented in humans [14] shows that the number of rhythmic genes in the liver is very low as compared to other tissues such as visceral adipose tissue or tissues in the heart, while in mice, the liver has the highest number of rhythmic genes [1]. In addition to species differences, other differences that may explain different results in the literature may relate to genetic heterogeneity, environmental or behavioral factors. As described, our work is more similar to that from [14], in showing that the liver has fewer rhythmic genes in humans as compared to in mice [1].

CIRCUST methodology provides a new insight with regard to the inherent presence of variability in core clock genes’ peak phase locations across tissues. This latter point evinces that there is no reason for considering the average of peak expressions for the analysis of clock molecular networks as done by [14]. Despite the heterogeneity between tissues shown by CIRCUST, tissue-specific molecular clock networks’ analyses show peak phases’ of TOP gene expressions clustered around dawn and dusk, with a quiescent period in-between as usually happens for diurnal primates as explained in [24].

CIRCUST presents several limitations to be considered. The first is regarding the assumption that the relative phase angle between the two selected clock genes is maintained across tissues and between people. We address this limitation by testing a modified model built on an alternative second clock gene of choice (*Dbp* primary and *Cry2* secondary). In addition, CIRCUST can derive the sequence and directionality of the temporal order of samples derived from a single-sample database to a high degree, but it does not make any assumptions or predictions regarding the phase angle between the circadian clock gene rhythms and local clock time. Finally, a limitation in this case of the GTEx dataset is that its population is heterogeneous in many ways including disease state, medication use, and environmental exposures. In addition, most tissues are heterogeneous with respect to cell type composition, and different cell types may have different properties regarding rhythmicity [42]. Even in a homogeneous tissue, individual cells may not be synchronized.

CIRCUST represents a step forward towards the building of a daily rhythm gene expression atlas in humans. Among the future directions, further systematic head-to-head comparisons of CIRCUST with other analytical methods are needed to determine the relative performance of each method under different conditions and in different populations and species. In particular, the study of covariates in the GTEx dataset to develop atlas comparisons regarding different demographic or clinical populations and conditions is important. Also, there is a need for future analysis in larger datasets because seasonality may interact with geographic location, the season could be an important covariate. This may be especially relevant for the sun-exposed skin from lower leg tissue. Finally, pathway analyses to follow up on the hits derived from our CIRCUST analyses are needed to advance understanding of tissue-specific and across-tissue rhythmic biological processes.

## Supporting information

Daily rhythm gene expression atlas in humans

## Data and Code availability

CIRCUST methodology is publicly available at https://github.com/yolandalago/CIRCUST/.

## Supplemental Materials

Supplemental material for this paper is available online.

## Author Contribution

Y.L., C.R., R.S. and F.A.J.L.S conceived of the presented idea. R.S. provided partial data. Y.L. and C.R. developed the theoretical proposal, and conceptual design and analyzed the results. Y.L. developed computational code, processed the data, performed the computations, and designed the analysis to validate the methodology. Y.L. wrote the manuscript with input from all authors. All authors approved the final manuscript.

