## Supplementary Materials for "CIRCUST: a novel methodology for temporal order reconstruction of molecular rhythms; validation and application towards a daily rhythm gene expression atlas in humans"

This supplementary material appends several issues mentioned in the manuscript. First, supplemental Figures and Tables are provided in order to thoroughly describe the methods and results given in the main text. Next, a supplemental text is included that is devoted to expanding several methodological details mentioned in the paper.

### 1 Supplementary Figures

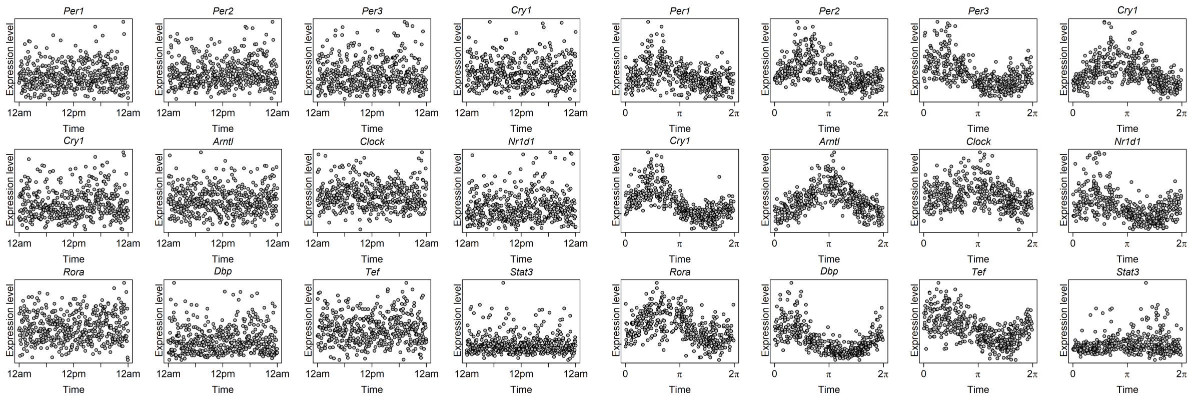

Figure S1: Core clock gene expression patterns from Skin sun-exposed (Lower leg) from GTEx dataset. Left: gene expressions as a function of TOD times. Right: gene expressions as a function of CIRCUST estimated times ($[0, 2\pi)$).

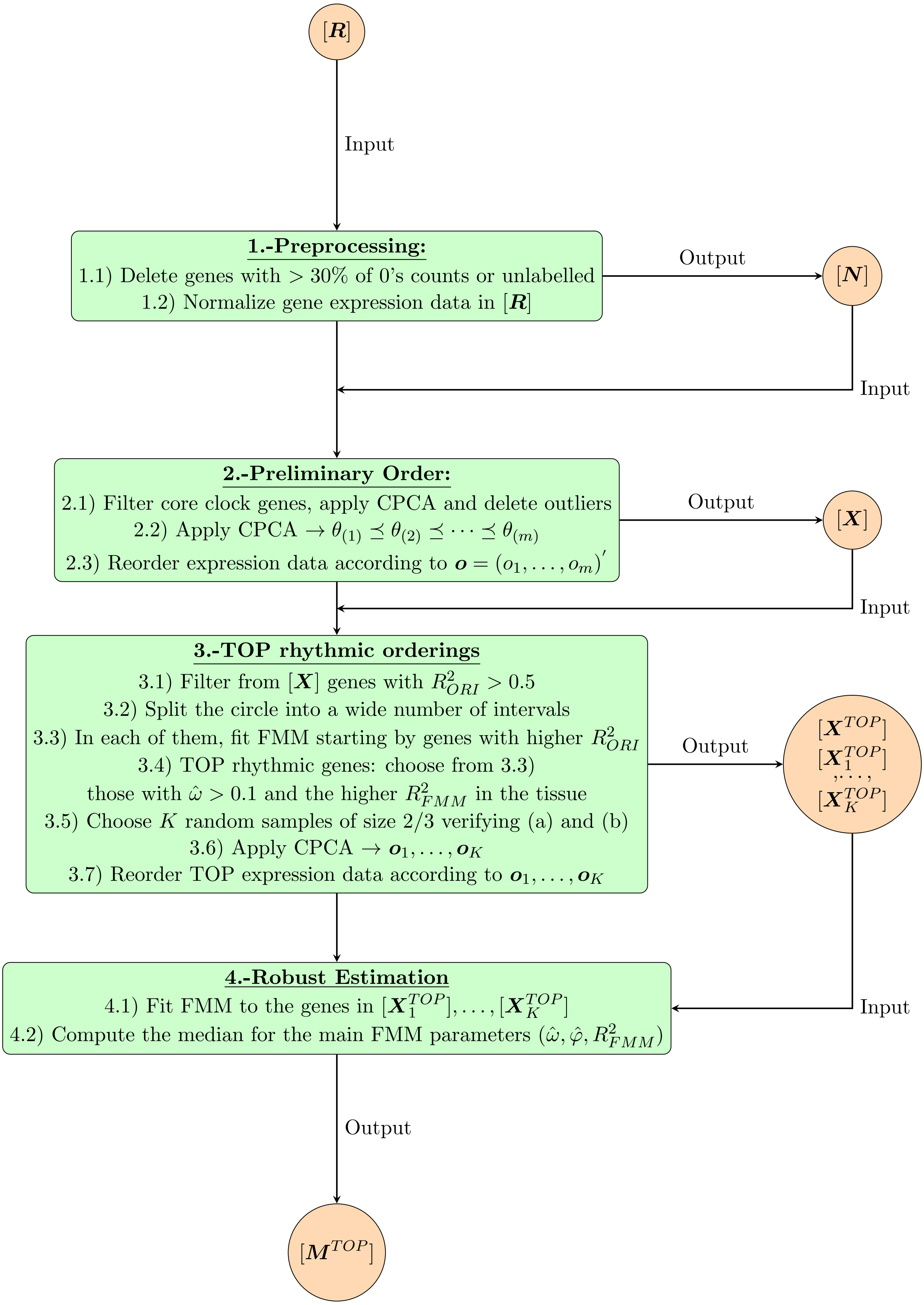

Figure S2: Outline of the CIRCUST methodology.

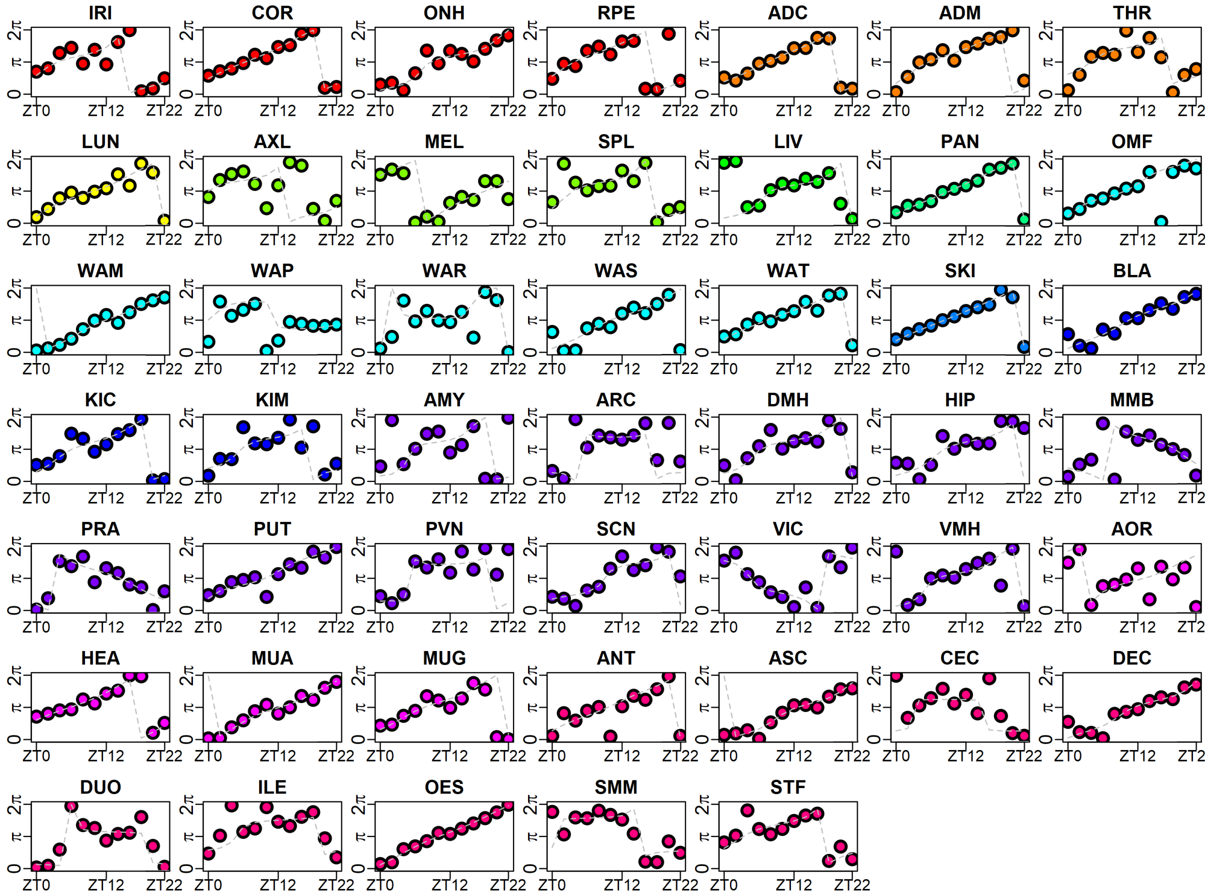

Figure S3: Circular-circular regression model for the real clock times (X-axis) and CIRCUST$_{prior}$ estimated times (Y-axis) across the 47 baboons' tissues selected. Horizontal axis: sampling real clock times along 24-h (ZT0,ZT2,...,ZT22). Vertical axis: CIRCUST estimated times in $[0, 2\pi)$. Time 0-h is the same as 24-h and the phase 0 is the same as $2\pi$. The diagonal line observed for most of the tissues is used as a marker of the coherence between the orders. Colors match with the 12 functional organs groups considered in GSE98965. See Table S1 for tissue names.

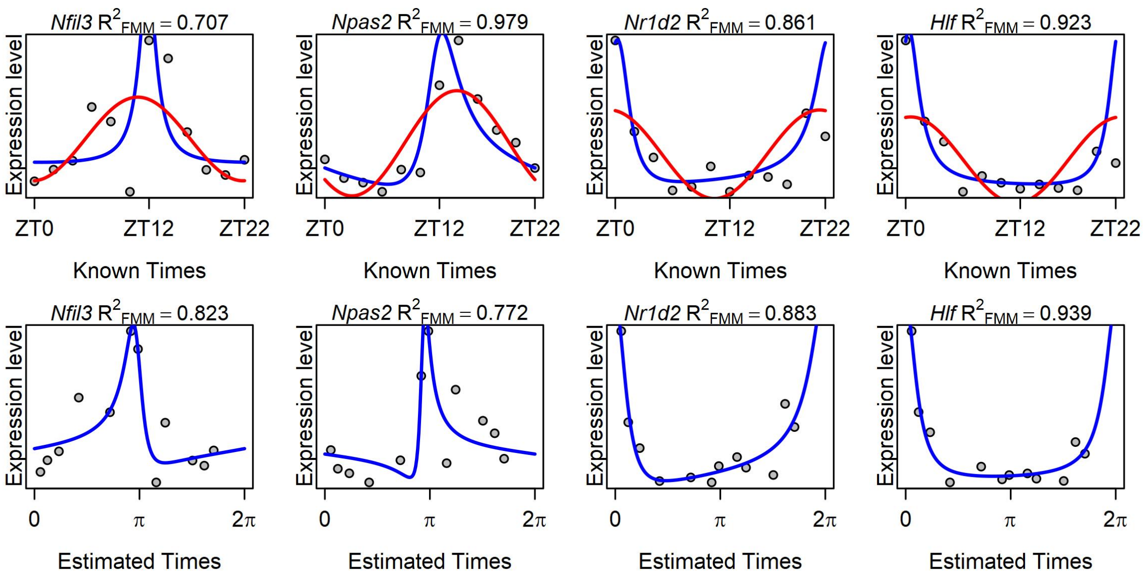

Figure S4: Expression of selected clock genes *Nfil3*, *Npas2*, *Nr1d2* and *Hlf* in White Adipose Mesenteric (WAM) tissue from baboons (GSE98965). Top panels: expressions as function of known times ZT0, ZT2,...,ZT22. Bottom panels: expressions as function of CIRCUST estimated times. FMM predictions are shown as blue solid lines. Cosinor predictions are shown as red solid lines.

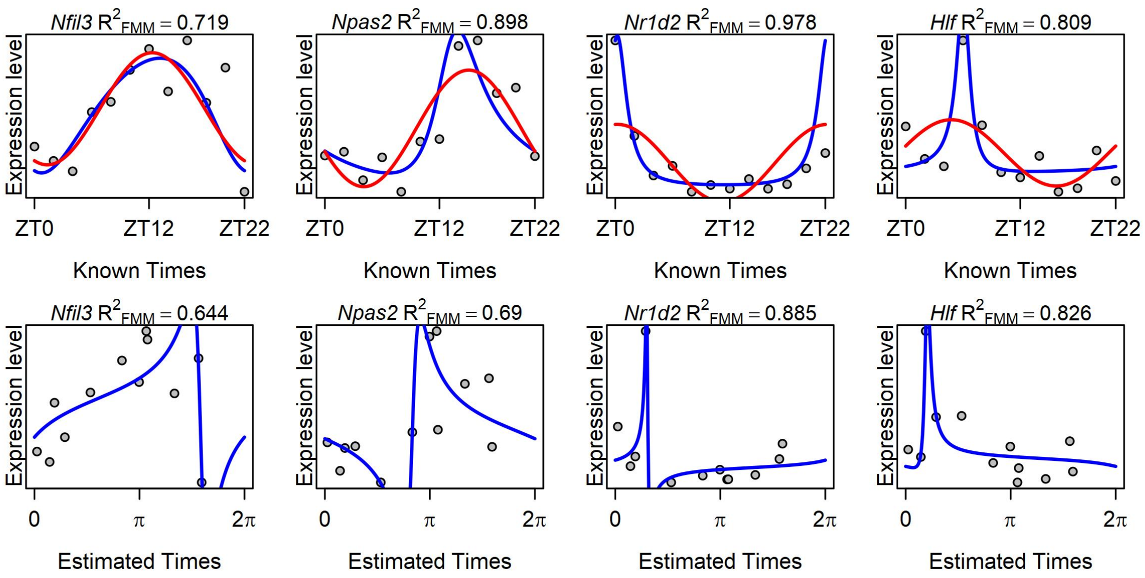

Figure S5: Expression of selected clock genes *Nfil3*, *Npas2*, *Nr1d2* and *Hlf* in Ascending Colon (ASC) tissue from baboons (GSE98965). Top panels: expressions as function of known times ZT0, ZT2,...,ZT22. Bottom panels: expressions as function of CIRCUST estimated times. FMM predictions are shown as blue solid lines. Cosinor predictions are shown as red solid lines.

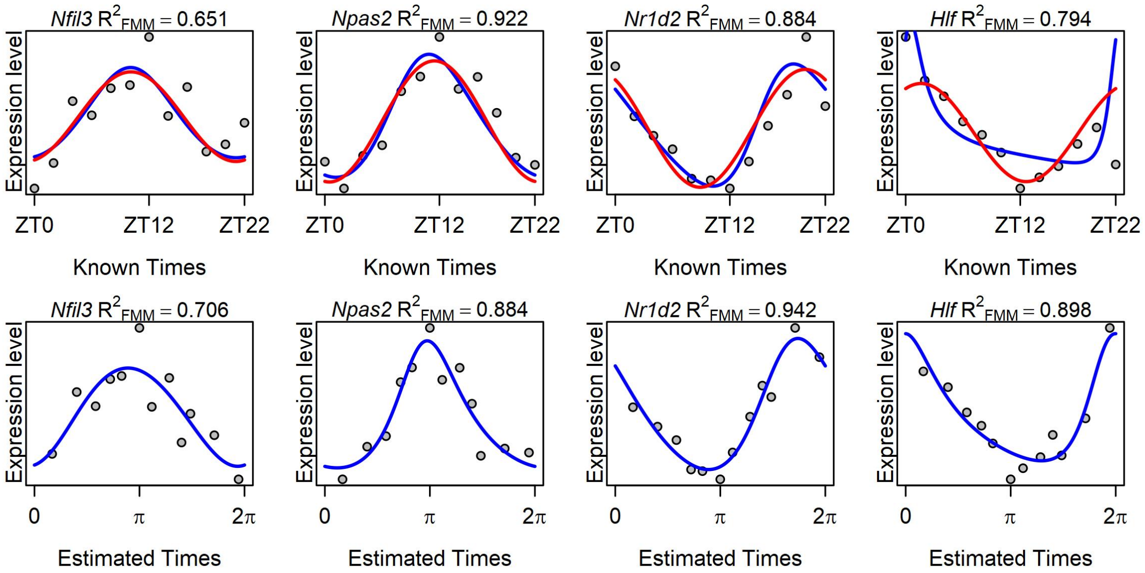

Figure S6: Expression of selected clock genes *Nfil3*, *Npas2*, *Nr1d2* and *Hlf* in Skin (SKI) tissue from baboons (GSE98965). Top panels: expressions as function of known times ZT0, ZT2,...,ZT22. Bottom panels: expressions as function of CIRCUST estimated times. FMM predictions are shown as blue solid lines. Cosinor predictions are shown as red solid lines.

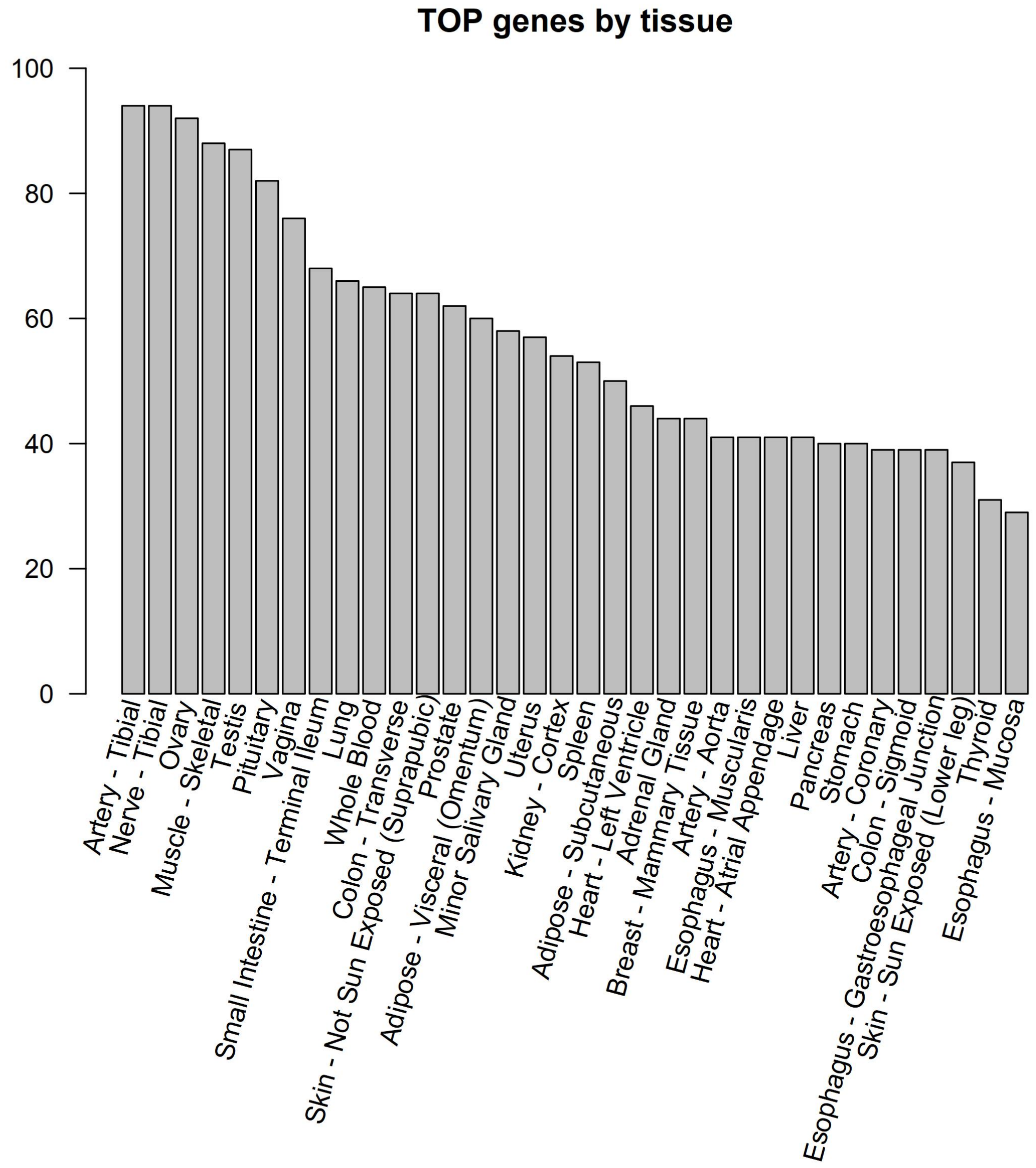

Figure S7: Number of TOP rhythmic genes across the GTEx. Tissues are shown in decreasing the number of TOP rhythmic genes.

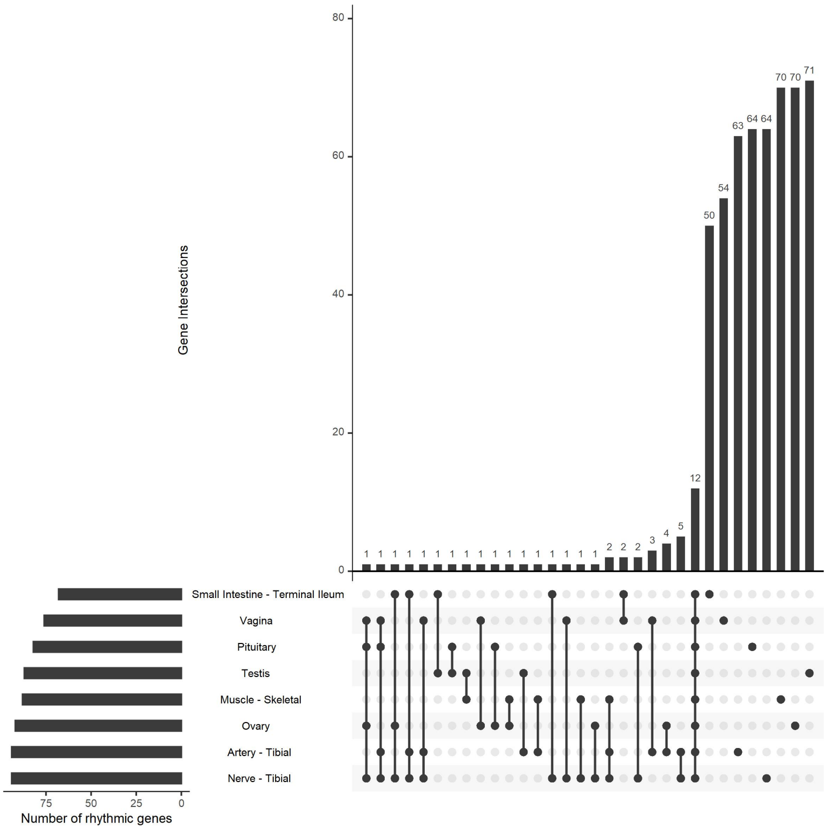

Figure S8: UpSetR [1] technique to visualize TOP rhythmic gene intersections across the 34 tissues. Dots denote single tissue intersections. Bars match tissues with common TOP rhythmic genes. For better illustration, only the eight tissues with the higher number of TOP genes have been included in the analysis.

### 2 Supplementary Tables

Table S1: Baboons tissue characterization. First column: Tissue abbreviation. Second column: Tissue name. Third column: $R^2_{Ave}$, average rhythmicity measure $R^2_{FMM}$ for the 12 core clock genes as a function of estimated times. Fourth column: $\rho$ goodness of fit measure fit measure for circular-circular regression between the real times (ZT0,ZT2,...,ZT22) and the CIRCUST estimated times ($[0, 2\pi)$). Tissues are restricted to those with $R^2_{Ave} > 0.7$ to guarantee the consistency of the results. Lines separate functional groups.

| Abbreviation | Tissue | $R^2_{Ave}$ | $\rho$ |
|---|---|---|---|
| IRI | Iris | 0.811 | 0.800 |
| COR | Cornea | 0.846 | 0.958 |
| ONH | Optic Nerve Head | 0.807 | 0.874 |
| RPE | Retina Pigment Epithelium | 0.740 | 0.871 |
| ADC | Adrenal Cortex | 0.787 | 0.947 |
| ADM | Adrenal Medulla | 0.799 | 0.899 |
| THR | Thyroid | 0.810 | 0.723 |
| LUN | Lung | 0.803 | 0.911 |
| AXL | Axilliary Lymphonodes | 0.724 | 0.615 |
| MEL | Mesenteric Lymphonodes | 0.746 | 0.833 |
| SPL | Spleen | 0.703 | 0.741 |
| LIV | Liver | 0.730 | 0.777 |
| PAN | Pancreas | 0.812 | 0.960 |
| OMF | Omental Fat | 0.796 | 0.892 |
| WAM | White Adipose Mesenteric | 0.814 | 0.964 |
| WAP | White Adipose Mesenteric | 0.745 | 0.609 |
| WAR | White Adipose Perirenal | 0.813 | 0.708 |
| WAS | White Adipose Subcutaneous | 0.811 | 0.878 |
| WAT | White Adipose Tissue | 0.892 | 0.961 |
| SKI | Skin | 0.788 | 0.953 |
| BLA | Bladder | 0.745 | 0.880 |
| KIC | Kidney Cortex | 0.811 | 0.765 |
| KIM | Kidney Medulla | 0.764 | 0.728 |
| AMY | Amygdala | 0.702 | 0.686 |
| ARC | Arcuate Nucleus | 0.745 | 0.680 |
| DMH | Dorsomedial Hypothalamus | 0.703 | 0.728 |
| HIP | Hippocampus | 0.704 | 0.734 |
| MMB | Mammilary Bodies | 0.735 | 0.789 |
| PRA | Preoptic Area | 0.702 | 0.741 |
| PUT | Putamen | 0.784 | 0.813 |
| PVN | Paraventricular Nuclei | 0.728 | 0.744 |
| SCN | Suprachiasmatic Nuclei | 0.711 | 0.729 |
| VIC | Visual Cortex | 0.740 | 0.706 |
| VMH | Ventromedial Hypothalamus | 0.776 | 0.806 |
| AOR | Aorta | 0.742 | 0.660 |
| HEA | Heart | 0.841 | 0.935 |
| MUA | Muscle Abdominal | 0.799 | 0.953 |
| MUG | Muscle Gastrocnemian | 0.701 | 0.884 |
| ANT | Antrum | 0.835 | 0.740 |
| ASC | Ascending Colon | 0.746 | 0.959 |
| CEC | Cecum | 0.701 | 0.734 |
| DEC | Descending Colon | 0.739 | 0.846 |
| DUO | Duodenum | 0.816 | 0.747 |
| ILE | Ileon | 0.709 | 0.726 |
| OES | Oesophagus | 0.903 | 0.973 |
| SMM | Smooth Muscle | 0.741 | 0.674 |
| STF | Stomach Fundus | 0.824 | 0.734 |

Table S2: GTEx donor distribution by sex, age, and cause of death. Death was classified as follows. Fast: death due to accident, blunt force trauma, or suicide; Intermediate: patients who were ill but death was unexpected; Slow: death after a long illness; Sudden-Natural: fast death of natural causes, sudden unexpected deaths; Ventilator: all cases on a ventilator immediately before death.

| | | | | | | |
|---|---|---|---|---|---|---|
| **Sex** | Male | Female | | | | |
| | 65.86% | 34.14% | | | | |
| **Age** | 20-29 | 30-39 | 40-49 | 50-59 | 60-69 | 70-79 |
| | 8.05% | 7.25% | 15.94% | 32.85% | 32.69% | 3.22% |
| **Death** | Fast | Intermediate | Slow | Sudden-Natural | Ventilator | |
| | 4.35% | 4.99% | 12.56% | 26.73% | 50.72% | |

Table S3: $R^2_{FMM}$ for the time course expression of the 12 core clock genes as a function of CIRCUST times across the 34 GTEx tissues analyzed. Only the peaks of the core clock genes with $R_F^2MM > 0.3$ are shown in Figure 7.

| | *Per1* | *Per2* | *Per3* | *Cry1* | *Cry2* | *Arntl* | *Clock* | *Nr1d1* | *Rora* | *Dbp* | *Tef* | *Stat3* |
|---|---|---|---|---|---|---|---|---|---|---|---|---|
| Adipose - Subcutaneous | 0.184 | 0.466 | 0.504 | 0.317 | 0.254 | 0.393 | 0.31 | 0.231 | 0.52 | 0.586 | 0.178 | 0.353 |
| Adipose - Visceral (Omentum) | 0.273 | 0.217 | 0.372 | 0.571 | 0.191 | 0.088 | 0.34 | 0.359 | 0.508 | 0.455 | 0.397 | 0.671 |
| Adrenal Gland | 0.109 | 0.074 | 0.531 | 0.308 | 0.176 | 0.366 | 0.559 | 0.396 | 0.34 | 0.714 | 0.661 | 0.446 |
| Artery - Aorta | 0.36 | 0.276 | 0.613 | 0.191 | 0.636 | 0.499 | 0.25 | 0.316 | 0.19 | 0.527 | 0.559 | 0.147 |
| Artery - Coronary | 0.155 | 0.36 | 0.666 | 0.365 | 0.491 | 0.565 | 0.496 | 0.229 | 0.514 | 0.568 | 0.526 | 0.339 |
| Artery - Tibial | 0.338 | 0.281 | 0.424 | 0.039 | 0.379 | 0.488 | 0.306 | 0.046 | 0.176 | 0.359 | 0.396 | 0.178 |
| Breast - Mammary Tissue | 0.405 | 0.54 | 0.687 | 0.445 | 0.419 | 0.637 | 0.241 | 0.517 | 0.613 | 0.61 | 0.324 | 0.22 |
| Colon - Sigmoid | 0.215 | 0.294 | 0.678 | 0.23 | 0.449 | 0.545 | 0.585 | 0.204 | 0.426 | 0.484 | 0.598 | 0.235 |
| Colon - Transverse | 0.535 | 0.077 | 0.592 | 0.517 | 0.652 | 0.367 | 0.671 | 0.377 | 0.75 | 0.332 | 0.422 | 0.299 |
| Esophagus - Gastroesophageal | 0.144 | 0.258 | 0.657 | 0.439 | 0.537 | 0.531 | 0.567 | 0.175 | 0.497 | 0.454 | 0.668 | 0.214 |
| Esophagus - Mucosa | 0.253 | 0.198 | 0.682 | 0.338 | 0.323 | 0.605 | 0.327 | 0.271 | 0.285 | 0.513 | 0.713 | 0.097 |
| Esophagus - Muscularis | 0.167 | 0.295 | 0.718 | 0.385 | 0.452 | 0.553 | 0.503 | 0.253 | 0.568 | 0.504 | 0.661 | 0.18 |
| Heart - Atrial Appendage | 0.338 | 0.306 | 0.64 | 0.437 | 0.373 | 0.342 | 0.303 | 0.211 | 0.48 | 0.451 | 0.668 | 0.345 |
| Heart - Left Ventricle | 0.222 | 0.104 | 0.692 | 0.557 | 0.481 | 0.564 | 0.714 | 0.227 | 0.376 | 0.471 | 0.758 | 0.341 |
| Kidney - Cortex | 0.478 | 0.822 | 0.754 | 0.75 | 0.723 | 0.75 | 0.8 | 0.395 | 0.675 | 0.794 | 0.767 | 0.877 |
| Liver | 0.54 | 0.232 | 0.485 | 0.239 | 0.559 | 0.156 | 0.561 | 0.316 | 0.555 | 0.497 | 0.142 | 0.613 |
| Lung | 0.358 | 0.269 | 0.236 | 0.559 | 0.456 | 0.203 | 0.03 | 0.278 | 0.442 | 0.301 | 0.549 | 0.477 |
| Minor Salivary Gland | 0.343 | 0.181 | 0.475 | 0.182 | 0.487 | 0.434 | 0.511 | 0.43 | 0.384 | 0.341 | 0.585 | 0.144 |
| Muscle - Skeletal | 0.442 | 0.302 | 0.451 | 0.338 | 0.409 | 0.557 | 0.559 | 0.166 | 0.507 | 0.42 | 0.287 | 0.199 |
| Nerve - Tibial | 0.157 | 0.451 | 0.583 | 0.311 | 0.226 | 0.368 | 0.299 | 0.155 | 0.399 | 0.6 | 0.255 | 0.213 |
| Ovary | 0.559 | 0.174 | 0.514 | 0.29 | 0.483 | 0.157 | 0.126 | 0.304 | 0.341 | 0.587 | 0.454 | 0.396 |
| Pancreas | 0.11 | 0.464 | 0.681 | 0.551 | 0.395 | 0.605 | 0.418 | 0.134 | 0.491 | 0.504 | 0.675 | 0.484 |
| Pituitary | 0.568 | 0.434 | 0.703 | 0.207 | 0.511 | 0.178 | 0.282 | 0.114 | 0.071 | 0.59 | 0.629 | 0.41 |
| Prostate | 0.101 | 0.167 | 0.491 | 0.309 | 0.286 | 0.283 | 0.49 | 0.22 | 0.516 | 0.473 | 0.472 | 0.433 |
| Skin - Not Sun Exposed | 0.415 | 0.235 | 0.519 | 0.182 | 0.506 | 0.471 | 0.175 | 0.46 | 0.547 | 0.564 | 0.176 | 0.133 |
| Skin - Sun Exposed | 0.193 | 0.313 | 0.543 | 0.251 | 0.509 | 0.503 | 0.216 | 0.308 | 0.402 | 0.605 | 0.463 | 0.05 |
| Small Intestine - Terminal Ile | 0.551 | 0.331 | 0.453 | 0.499 | 0.678 | 0.265 | 0.379 | 0.404 | 0.468 | 0.12 | 0.227 | 0.221 |
| Spleen | 0.354 | 0.338 | 0.351 | 0.334 | 0.458 | 0.46 | 0.641 | 0.309 | 0.495 | 0.108 | 0.393 | 0.587 |
| Stomach | 0.513 | 0.112 | 0.607 | 0.415 | 0.576 | 0.543 | 0.545 | 0.216 | 0.583 | 0.635 | 0.697 | 0.582 |
| Testis | 0.196 | 0.246 | 0.555 | 0.655 | 0.119 | 0.473 | 0.752 | 0.315 | 0.295 | 0.382 | 0.314 | 0.522 |
| Thyroid | 0.275 | 0.535 | 0.708 | 0.351 | 0.532 | 0.54 | 0.403 | 0.198 | 0.469 | 0.537 | 0.664 | 0.176 |
| Uterus | 0.235 | 0.349 | 0.456 | 0.341 | 0.594 | 0.313 | 0.196 | 0.457 | 0.561 | 0.559 | 0.443 | 0.432 |
| Vagina | 0.306 | 0.365 | 0.537 | 0.436 | 0.6 | 0.296 | 0.205 | 0.481 | 0.384 | 0.544 | 0.363 | 0.164 |
| Whole Blood | 0.499 | 0.593 | 0.556 | 0.673 | 0.753 | 0.693 | 0.721 | 0.597 | 0.682 | 0.462 | 0.488 | 0.509 |

Table S4: Estimated $t_U$ for the time course expression of the 12 core clock genes as a function of CIRCUST times across the 34 GTEx tissues analyzed. Only the peaks of the core clock genes with $R^2_F MM > 0.3$ are shown in Figure 7.

| | *Per1* | *Per2* | *Per3* | *Cry1* | *Cry2* | *Arntl* | *Clock* | *Nr1d1* | *Rora* | *Dbp* | *Tef* | *Stat3* |
|---|---|---|---|---|---|---|---|---|---|---|---|---|
| Adipose - Subcutaneous | 1.196 | 0.865 | 0.173 | 1.417 | 0.397 | 3.142 | 2.196 | 0.306 | 1.395 | 6.073 | 0.093 | 2.031 |
| Adipose - Visceral (Omentum) | 2.855 | 4.046 | 1.941 | 3.491 | 2.195 | 3.142 | 3.484 | 2.127 | 3.082 | 1.855 | 1.927 | 3.788 |
| Adrenal Gland | 5.408 | 0.95 | 0.806 | 2.683 | 3.553 | 3.142 | 2.214 | 0.627 | 1.519 | 0.533 | 0.805 | 3.105 |
| Artery - Aorta | 1.089 | 0.991 | 0.666 | 3.005 | 1.153 | 3.142 | 3.328 | 1.141 | 3.049 | 0.688 | 0.289 | 3.341 |
| Artery - Coronary | 1.581 | 1.888 | 0.786 | 2.421 | 1.607 | 3.142 | 2.823 | 1.207 | 2.742 | 0.67 | 0.768 | 2.645 |
| Artery - Tibial | 2.124 | 1.719 | 1.352 | 4.382 | 1.717 | 3.142 | 2.795 | 1.971 | 2.637 | 0.549 | 1.912 | 4.578 |
| Breast - Mammary Tissue | 0.755 | 0.557 | 0.194 | 1.444 | 0.335 | 3.142 | 2.137 | 1.072 | 2.159 | 0.355 | 0.137 | 1.66 |
| Colon - Sigmoid | 1.412 | 0.801 | 0.889 | 3.654 | 0.796 | 3.142 | 3.01 | 0.786 | 2.149 | 0.497 | 1.038 | 3.897 |
| Colon - Transverse | 2.655 | 2.429 | 2.86 | 3.096 | 2.819 | 3.142 | 3.164 | 2.876 | 3.014 | 2.818 | 2.896 | 2.802 |
| Esophagus - Gastroesophageal | 1.332 | 0.831 | 0.82 | 3 | 0.964 | 3.142 | 2.388 | 0.821 | 1.758 | 0.458 | 0.814 | 2.549 |
| Esophagus - Mucosa | 1.77 | 1.546 | 0.804 | 3.521 | 1.298 | 3.142 | 2.919 | 1.271 | 2.125 | 0.881 | 0.867 | 4.629 |
| Esophagus - Muscularis | 1.8 | 1.245 | 1.031 | 3.291 | 1.188 | 3.142 | 2.729 | 0.48 | 2.322 | 0.547 | 0.869 | 3.574 |
| Heart - Atrial Appendage | 2.261 | 1.907 | 1.207 | 3.01 | 2.229 | 3.142 | 2.342 | 2.206 | 2.522 | 1.002 | 1.05 | 3.113 |
| Heart - Left Ventricle | 0.213 | 0.898 | 0.892 | 2.993 | 1.23 | 3.142 | 2.512 | 0.142 | 2.172 | 0.602 | 1.064 | 3.301 |
| Kidney - Cortex | 2.697 | 2.266 | 2.145 | 2.485 | 2.288 | 3.142 | 2.431 | 2.587 | 2.519 | 1.83 | 2.21 | 2.649 |
| Liver | 2.695 | 2.154 | 2.181 | 3.868 | 2.2 | 3.142 | 4.369 | 2.511 | 3.338 | 2.141 | 2.296 | 4.696 |
| Lung | 2.123 | 2.763 | 1.928 | 3.005 | 2.547 | 3.142 | 2.525 | 2.382 | 2.922 | 1.809 | 1.325 | 4.007 |
| Minor Salivary Gland | 0.139 | 5.065 | 0.749 | 2.589 | 0.252 | 3.142 | 2.474 | 0.385 | 1.724 | 6.124 | 0.688 | 3.274 |
| Muscle - Skeletal | 2.12 | 1.82 | 1.811 | 2.763 | 1.941 | 3.142 | 2.903 | 2.113 | 2.499 | 0.599 | 1.645 | 2.629 |
| Nerve - Tibial | 1.368 | 1.591 | 1.036 | 2.437 | 1.473 | 3.142 | 2.199 | 5.588 | 1.967 | 0.616 | 0.87 | 2.718 |
| Ovary | 3.567 | 3.134 | 2.382 | 4.93 | 3.509 | 3.142 | 5.777 | 2.664 | 3.53 | 1.925 | 2.225 | 4.995 |
| Pancreas | 2.552 | 1.822 | 1.022 | 3.1 | 2.144 | 3.142 | 2.243 | 1.544 | 2.573 | 0.736 | 1.403 | 2.743 |
| Pituitary | 0.37 | 1.039 | 1.346 | 5.416 | 0.338 | 3.142 | 2.634 | 0.591 | 3.058 | 1.379 | 1.522 | 5.317 |
| Prostate | 2.952 | 3.8 | 2.106 | 4.058 | 2.097 | 3.142 | 3.158 | 2.435 | 2.932 | 1.832 | 1.861 | 4.192 |
| Skin - Not Sun Exposed | 2.193 | 3.29 | 1.069 | 2.38 | 1.882 | 3.142 | 1.258 | 2.115 | 2.57 | 1.409 | 1.305 | 5.077 |
| Skin - Sun Exposed | 1.621 | 1.742 | 1.09 | 2.284 | 1.458 | 3.142 | 2.927 | 1.202 | 1.929 | 0.73 | 0.811 | 3.645 |
| Small Intestine - Terminal Ileum | 2.973 | 2.884 | 2.611 | 2.815 | 2.924 | 3.142 | 2.9 | 1.762 | 2.961 | 1.652 | 2.631 | 2.759 |
| Spleen | 2.786 | 2.771 | 2.403 | 2.68 | 3.061 | 3.142 | 2.971 | 2.24 | 2.999 | 3.231 | 2.367 | 2.827 |
| Stomach | 2.935 | 4.583 | 1.964 | 3.745 | 2.645 | 3.142 | 3.277 | 1.922 | 2.994 | 1.643 | 2.221 | 3.559 |
| Testis | 2.917 | 3.859 | 2.682 | 4.261 | 3.673 | 3.142 | 4.044 | 2.451 | 3.072 | 2.248 | 2.89 | 2.466 |
| Thyroid | 0.914 | 0.978 | 1.012 | 2.768 | 1.118 | 3.142 | 2.557 | 1.216 | 2.641 | 0.829 | 0.915 | 2.223 |
| Uterus | 2.54 | 1.638 | 1.361 | 2.807 | 1.861 | 3.142 | 3.885 | 2.093 | 2.83 | 1.284 | 1.288 | 3.368 |
| Vagina | 3.012 | 0.851 | 1.078 | 1.707 | 1.649 | 3.142 | 5.757 | 1.712 | 1.535 | 0.961 | 1.439 | 2.845 |
| Whole Blood | 1.687 | 2.143 | 2.255 | 2.017 | 1.998 | 3.142 | 2.817 | 1.778 | 1.962 | 1.496 | 2.74 | 3.146 |

### 3 Supplemental text: Advanced Methodological details

#### 3.1 FMM model approach

The Frequency Modulated Möbius (FMM) proposed in [2] is a multi-purpose approach that combines a physically meaningful formulation with excellent statistical and computational properties. It is formulated as a signal plus error model where the signal is described parametrically as an oscillatory signal. The parametric formulation facilitates the interpretability and the derivation of essential elements in the analysis of oscillations. A distinguishing feature of the FMM is that it is formulated in terms of the phase ($\varphi$), which is an angular variable representing the periodic oscillation movement. All the methodological details that justify the mathematical formulation of the FMM models are given in [2].

Let assume that the time points are in $[0, 2\pi)$. In any other case, transform the time points $t' \in [t_0, T + t_0]$ by $t = \frac{(t'-t_0)2\pi}{T}$. The FMM signal plus error model is defined as follows.

**Definition 1.** FMM model.

For the observations $t_1 < \ldots < t_m$,

$$X(t_i) = \mu(t_i) + e(t_i) = M + A\cos(\varphi(t_i)) + e(t_i); \quad i = 1, \ldots, m$$

- $M \in \mathcal{R}, A \in \mathcal{R}^+$.
- $\varphi(t) = \beta + 2\arctan(\omega \tan(\frac{t-\alpha}{2})); \alpha, \beta \in [0, 2\pi), \omega \in [0, 1]$.
- $(e(t_1), \ldots, e(t_m))' \sim N_m(0, \sigma^2)$.

The FMM parameters characterize various aspects of a rhythmic pattern. The parameter $M$ is an intercept and $A$ measures the signal's amplitude. $\alpha$ is a phase location parameter, while $\omega$ and $\beta$ are parameters that describe the shape. Specifically, $\omega$ measures the sharpness, and $\beta$ measures skewness and indicates upward and/or downward peak direction. Note that a sinusoidal curve corresponds to $\omega = 1$.

Other important parameters of practical use are peak and trough times, denoted by $t_U$ and $t_L$, respectively, which are defined as follows:

$$t_U = \alpha + 2\arctan(\frac{1}{\omega}\tan(\frac{-\beta}{2}))$$

$$t_L = \alpha + 2\arctan(\frac{1}{\omega}\tan(\frac{\pi - \beta}{2}))$$

#### 3.2 CPCA Temporal order estimation. Starting point and direction choice

The solution of the temporal order estimation problem proposed in this work not only solves the mathematical problem of identifying a circular order, but also states the starting point and the clockwise or counterclockwise direction. Methodological details for these purposes are given below.

Let $[\boldsymbol{X}]$ be a gene expression matrix, and let $\boldsymbol{E}_1$ and $\boldsymbol{E}_2$ be the two first *eigengenes* of $[\boldsymbol{X}]$. Eigengenes are linear combination of the gene expressions along the direction of the most variation in the data [3]. Under rhythmicity, the mapping of $\boldsymbol{E}_1$ against $\boldsymbol{E}_2$ reveals an underlying circular structure, see panels (a), (b) and (c) in Figure S9.

The Circular Principal Component Analysis (CPCA) [4] is defined as the transformation in which the eigengenes are projected onto the unit circle as follows:

$$(e_{1,i}, e_{2,i}) = (\frac{E_{1,i}}{\sqrt{E_{1,i}^2 + E_{2,i}^2}}, \frac{E_{2,i}}{\sqrt{E_{1,i}^2 + E_{2,i}^2}}), i = 1, \ldots, m$$

From these projections, CPCA computes $\boldsymbol{\theta} = (\theta_1, \ldots, \theta_m)'$, the vector of angular phases that represents the temporal position of samples onto the unit circle ($[0, 2\pi)$), where $\theta_i = \arctan(\frac{e_{1,i}}{e_{2,i}}) \forall i = 1, \ldots, m$. The increasing order of these phases $\theta_{(1)} \prec \theta_{(2)} \prec \cdots \prec \theta_{(m)}$ induces a circular order $\mathbf{o}$ on data collection, see panels (d), (e) and (f) in Figure S9.

Each circular order represents $2m$ sampling time configurations depending on the choice of starting point and the direction. To make this choice, CIRCUST relies on three standard assumptions: (1) The peak phase of *Arntl* is set at $\pi$ which induces a starting point and facilitates comparisons. Moreover, as it is well-known that *Arntl* peaks in anticipation of the inactive period in mammals [5], then in this work $[0, \pi)$ and $[\pi, 2\pi)$ represent inferred light and dark periods, respectively; (2) *Dbp* peaks after ROR-phased genes (*Arntl, Npas, Clock*) [6, 7, 8]; (3) Peaks phases for the most of the core clock genes occur while the active period $[0, \pi)$ [9, 10]. These two latter assumptions define clockwise or counterclockwise direction.

In CIRCUST$_{prior}$ the assumption (2) can be refined, in terms of peak phases' order restrictions, incorporating additional knowledge regarding the molecular clock network of the species or experiment.

#### 3.3 CPCA Outliers sample detection

The role of CPCA in this work is twofold. In addition to providing a solution for the temporal order estimation problem, it can be used, in conjunction with the FMM predictions, as an outlier identification tool. Consider the mapping of the two first eigengenes computed on the $[\boldsymbol{N}]$ from the 12 core clock genes. The observation pairs samples at a

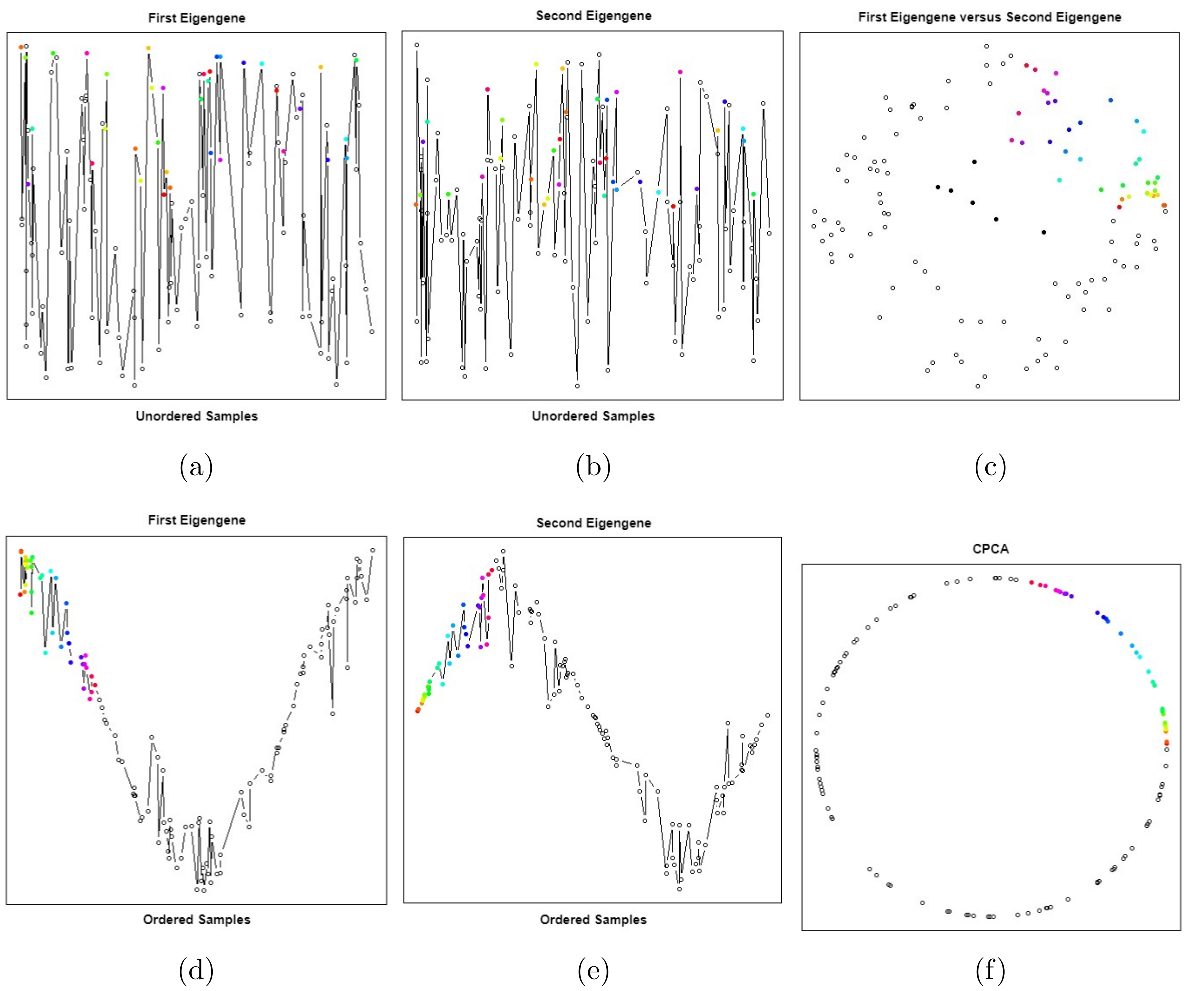

Figure S9: CPCA performance illustration. (a): First eigengene from the unordered expression matrix. (b): Second eigengene from the unordered expression matrix. (c): (a) versus (b) mapping. Black dots are outliers samples to be deleted, see Section 3.3 (d): First eigengene ordered regarding the increased order of the angular values given in $\boldsymbol{\theta}$. (e): Second eigengene ordered regarding the increased order of the angular values given in $\boldsymbol{\theta}$. (f) Eigengenes' projection onto the unit circle regarding the increased order of the angular values given in $\boldsymbol{\theta}$. Colors illustrate the order among the samples with the lower angular values in $\boldsymbol{\theta}$.

neighbourhood of the origin $(0, 0)$ disrupt the underlying circular structure of the data, see the inner black dots in Figure S9 (c). Let $L_{E_i}$ denote the distance in absolute value between $(0, 0)$ and the pairs $(E_{1,i}, E_{2,i})$, for $i = 1, \ldots, m$. Then, the FMM adequacy is assessed based on the standardized residuals $(r_i, i = 1, \ldots, n)$ of the 12 core clock genes. Paired samples violating $L_{E_i} < 0.1$ or $r_i > 3$ are declared as outliers and deleted from $[\boldsymbol{N}]$.

##### 3.4 $R^2-$based goodness of fit criteria

Given a rhythmicity model (ORI, FMM, Cosinor), this work employs a $R^2-$based goodness of fit criteria to assess rhythmicity following the lines given in [2, 11]. This measure is defined as follows:

$$R^2_{Model}(\boldsymbol{X}^g) = \frac{\sum_{i=1}^m (X_i^g - \hat{X}_i^g)^2}{\sum_{i=1}^m (X_i^g - \overline{X}^g)^2}, \tag{1}$$

where $\boldsymbol{X}^g = (X_1^g, \ldots, X_m^g)'$ denotes the gene expression of gene $g$, $\hat{\boldsymbol{X}}^g$ is the predicted expression pattern from the rhythmicity model and $\overline{X}^g$ denotes $\boldsymbol{X}^g$ average expression value.
